# Direct activation of RA signaling in cardiomyocytes protects hearts from apoptosis after myocardial infarction in mice

**DOI:** 10.1101/2020.12.04.395970

**Authors:** Fabio Da Silva, Fariba Jian Motamedi, Amelie Tison, Lahiru Chamara Weerasinghe Arachchige, Stephen T. Bradford, Jonathan Lefebvre, Pascal Dollé, Norbert B. Ghyselinck, Kay Dietrich Wagner, Andreas Schedl

## Abstract

Retinoic acid (RA) is an essential signaling molecule for cardiac development and plays a protective role in the heart after myocardial infarction (MI). In both cases, the effect of RA signaling on cardiomyocytes, the principle cell type of the heart, has been reported to be indirect. Here we have developed an inducible murine transgenic RA-reporter line using *CreER*^*T2*^ technology that permits lineage tracing of RA-responsive cells and faithfully recapitulates endogenous RA activity in multiple organs during embryonic development. Strikingly, we have observed a direct RA response in cardiomyocytes during mid-late gestation and after MI. Ablation of RA signaling through deletion of the *Aldh1a1/a2/a3* genes encoding RA-synthesizing enzymes leads to increased cardiomyocyte apoptosis in adults subjected to MI. RNA sequencing analysis reveals *Tgm2* and *Ace1*, two genes with well-established links to cardiac repair, as potential targets of RA signaling in primary cardiomyocytes, thereby providing novel links between the RA pathway and heart disease.

## INTRODUCTION

Retinoic acid (RA), the active derivative of vitamin A, plays essential roles in cell growth, differentiation, and organogenesis (*1*, *2*). RA is synthesized in two oxidative steps, with the second, rate-limiting step being carried out by three retinaldehyde dehydrogenases (ALDH1A1, ALDH1A2, ALDH1A3). Once synthesized, RA can activate or repress the transcription of various genes by binding to nuclear retinoic acid receptors (RARA, RARB, RARG), which form heterodimers with retinoid X receptors (RXRA, RXRB, RXRG) (*3*). Unlike most other embryonic signals that are peptidic in nature and act through membrane receptors, RA is a very small lipophilic molecule that cannot be easily detected by conventional means (*4*). Hence, determining the activity of RA signaling in specific cell types or tissues is challenging, and has traditionally relied on RA reporter lines.

In mice, several transgenic RA reporter lines have been developed in order to better understand the RA signaling pathway. These include the *Tg(RARE-Hspa1b/lacZ)12Jrt* (hereafter referred to as *RARE-LacZ*), *Tg(RARE/Hspa1b-cre)1Dll* (referred to as *RARE-Cre*) and *RARE-Luciferase* lines, among others (*5*, *6*, *7*). All three lines utilize multiple copies of the *Rarb* retinoic acid response element (RARE) to drive β-galactosidase (β-gal), Cre recombinase or Luciferase expression, and they have proven useful in understanding the wide range of RA responses during embryonic development and disease. Yet, due to inherent technical problems, these lines are limited in their ability to illustrate the full scope of RA activity in the mouse. For instance, the *RARE-LacZ* line is highly efficient at detecting acute RA activity at various stages of embryonic development, but is unable to trace the long-term fate of RA-responsive cells (*5*). Moreover, the β-gal protein is very stable and persists in a tissue long after its expression ceases, so measurement of its activity may not reflect RA signaling in real time. The *RARE-Cre* is also very effective at detecting RA-responses in tissues, but since it promotes constitutive Cre expression and permanent labeling of cells, the timing of the response cannot be determined (*6*). The *RARE-Luciferase* line can detect RA activity *in vivo*, but due to its use of Luciferase as a reporter, the cell types responding to RA in specific tissues cannot be identified by conventional immunohistochemistry (*7*). Hence, despite the wide variety of lines available, there remains a need for new tools capable of detecting RA activity in a controlled, reliable and efficient manner in order to better understand the roles of RA signaling in complex organs.

One such organ with which RA signaling is intimately linked is the heart. In order to meet high metabolic demands and ensure enough blood is efficiently pumped throughout the body, the mammalian heart must develop a compact layer of cardiomyocytes, whose coordinated contractions drive the pumping action of the heart (*8*). Myocardial compaction occurs between embryonic day (E)10-14 in the mouse, and involves the proliferation and differentiation of cardiomyoblasts, the precursors of cardiomyocytes (*9*, *10*, *11*). RA signaling is intricately linked with myocardial compaction and both *Rxra*-null and *Aldh1a2*-null mutant mice display severe hypoplasia of the compact layer (*12*, *13*, *14*). Despite having a pronounced effect on the myocardium, it has been suggested that the effects of RA signaling on developing cardiomyocytes is indirect. For instance, studies have shown that RA, synthesized by ALDH1A2 in the epicardium, works in an autocrine manner with RXRA functioning as the principle receptor (*9*, *11*). The ALDH1A2/RXRA axis stimulates the production of mitogens such as FGFs, which are then secreted to the myocardium to promote cell proliferation (*15*, *16*, *17*). It has also been suggested that ALDH1A2/RXRA signaling in the liver and placenta promote the production of erythropoietin (EPO) and distribution of glucose, respectively, both of which sequentially activate epicardial *Igf2* expression (*18*, *19*). IGF2 then stimulates myocardial proliferation and compaction. Hence, whether the effects of retinoids are within the epicardium or extra-cardiac tissues (or both), it appears that RA signaling does not act directly in cardiomyocytes.

In regard to mammalian heart disease, many studies conducted in adult rats and mice suggest that RA signaling stimulates cardiac repair after myocardial infarction. Vitamin A-deficient rats subjected to ligation of the left anterior descending artery (LAD), a model of MI, exhibit increased cardiomyocyte hypertrophy and interstitial collagen deposition, while supplementation of RA to mice following ischaemia/reperfusion surgery results in decreased apoptosis and smaller infarct zones (*20*, *21*). In both cases the cell types responding to RA treatment were not identified. Direct evidence that RA signaling is reactivated in the heart post MI comes from a study performed with the *RARE-Luciferase* reporter line. In this study, the authors detected RA signaling in damaged hearts 24 hours to one week post MI (*7*). Cardiac fibroblasts, and not cardiomyocytes, were determined to be the principle cell-types responding to RA, once again suggesting an indirect effect of RA on the myocardium.

Despite the majority of studies suggesting that cardiomyocytes are not the major RA-responsive cell types in the heart, minor, albeit important roles for RA signaling in the mammalian myocardium have been identified. In a study by Guleria et al. 2010, it was demonstrated that stimulation of the RA pathway in cultured neonatal cardiomyocytes subjected to high-glucose conditions led to decreased apoptosis. The protective effect was attributed to RA-induced modulation of the Renin-angiotensin system (RAS). More recently, cardiomyocyte-specific deletion of the RARA receptor in adult mice led to increased cardiomyocyte hypertrophy, excessive reactive oxygen species (ROS) accumulation and calcium mishandling defects (*23*). Furthermore, in non-mammalian vertebrates capable of naturally restoring damaged cardiac tissue without scar formation, it has been shown that RA signaling is indispensable for the regenerative response (*24*). For example, in adult zebrafish subjected to ventricular resection injuries, *Aldh1a2* mRNA was found to be upregulated in the endocardium and epicardium, and inhibition of RA signaling through transgenic expression of dominant-negative RARA led to drastically reduced cardiomyocyte proliferation (*25*). Altogether, these studies suggest RA signaling is active in cardiomyocytes, and that the RA pathway may be involved in protecting and/or repairing damaged heart muscle.

Here we have developed a novel RA reporter line using *CreER*^*T2*^ technology (*RARECreER*^*T2*^) that closely mimics well known RA reporter lines and allows for timed, cell type-specific tracing of RA-responsive cells. Using this line, we have detected a myocardial-specific response to RA during mid-late stages of development (E11-18), and have observed RA activity in cardiomyocytes of adult mice subjected to MI. In order to better understand the role of RA in cardiac repair, we have crossed *lox*P-flanked (floxed) alleles of the *Aldh1a* enzymes (*26*-*28*) with the ubiquitously expressed and inducible *Tg(CAG-cre/Esr1*)5Amc* line, hereafter referred to as *CAGGCreER*^*TM*^ (*29*), to generate time-controlled loss of functions. By temporally deleting the *Aldh1a* alleles in adult mice then subjecting them to MI, we have observed a drastic increase in cardiomyocyte-specific apoptosis. RNA sequencing of primary cardiomyocytes treated with RA followed by genome-wide analysis reveals RA can stimulate a notable transcriptional response in cardiomyocytes. Importantly, some of the identified RA-responsive genes, such as transglutaminase 2 (*Tgm2*) and angiotensin converting enzyme (*Ace1*), have well characterized roles in cardiac repair, thereby providing new molecular links between RA signaling and heart disease.

## RESULTS

### A novel RARECreER^T2^ line recapitulates endogenous RA signaling in developing mouse embryos

The RA signaling pathway is extremely dynamic, acting on various tissues at different stages of embryonic development (*1*, *3*). In order to better understand the timing and identity of the cell types responding to RA, we developed a novel tamoxifen-inducible RA reporter line using *CreER*^*T2*^ technology (Fig. 1A, schematic). Previous work has shown that RA signaling, activated by expression of *Aldh1a2* in caudal regions of the embryo, is first detected around E7.5 (*5*, *30*). To see if we could detect this early RA activity, we crossed our line with the *Gt(ROSA)26Sor*^*tm1Sor*^ reporter line (referred to as *R26L*) (*31*) and administered tamoxifen (TAM) one day earlier, at E6.5, to account for tamoxifen processing and eventual reporter activation (Fig. 1A). Strikingly, analysis of embryos at E9.5 via whole-mount X-gal staining demonstrated high reporter activity up to posterior hindbrain/pharyngeal regions of the embryo (Fig. 1B, left panel). Administration of tamoxifen at E7.5 followed by analysis at E10.5 revealed a similar pattern of staining, as well as efficient labeling of the developing limbs (Fig. 1B, right panel). The X-gal staining at both time-points closely matched the staining pattern observed with other RARE reporter lines such as the *RARE-LacZ* and *RARE-Cre*, and is consistent with the conserved roles of RA signaling role in embryo posteriorization, and limb bud formation (*5*, *6*, *32*-*35*). However, we detected very few X-gal^+^ cells in the forebrains of our *RARECreER*^*T2*^ embryos, a known site of RA activity (*5*, *6*, *36*). To address this issue, we crossed our line with the *Gt(ROSA)26Sor*^*tm4(ACTB-tdTomato,- EGFP)Luo*^ reporter line (referred to as *mTmG*) (*37*), which expresses GFP protein upon Cre recombination and is more sensitive than the *R26L* line. Whole mount GFP immunofluorescence (IF) of embryos induced at E7.5 and sacrificed at E10.5 revealed efficient labeling of the forebrain (Fig. 1C; Supplementary movie 1). To examine the full spectrum of the RA response detected with the *RARECreER*^*T2*^ line, we repeated the lineage tracing experiments with two pulses of tamoxifen at E7.5 and E8.5 to improve the overall recombination efficiency, and then analyzed embryos at E11.5, when most organs are further developed. Analysis of whole embryo sections by GFP IF revealed efficient and consistent labeling of various well-known RA-responsive organs such as the lungs (*38*), liver (*18*), spinal cord (*39*), heart (*40, 41*) and differentiating somites (*5*, *42*-*44*) in all embryos analyzed (Fig. 1D). Somite-specific labeling, as revealed by co-IF for MyoD and GFP, was also detected with a single pulse at E8.5 followed by analysis at E11.5 (Fig. 1E).

**Fig. 1:**
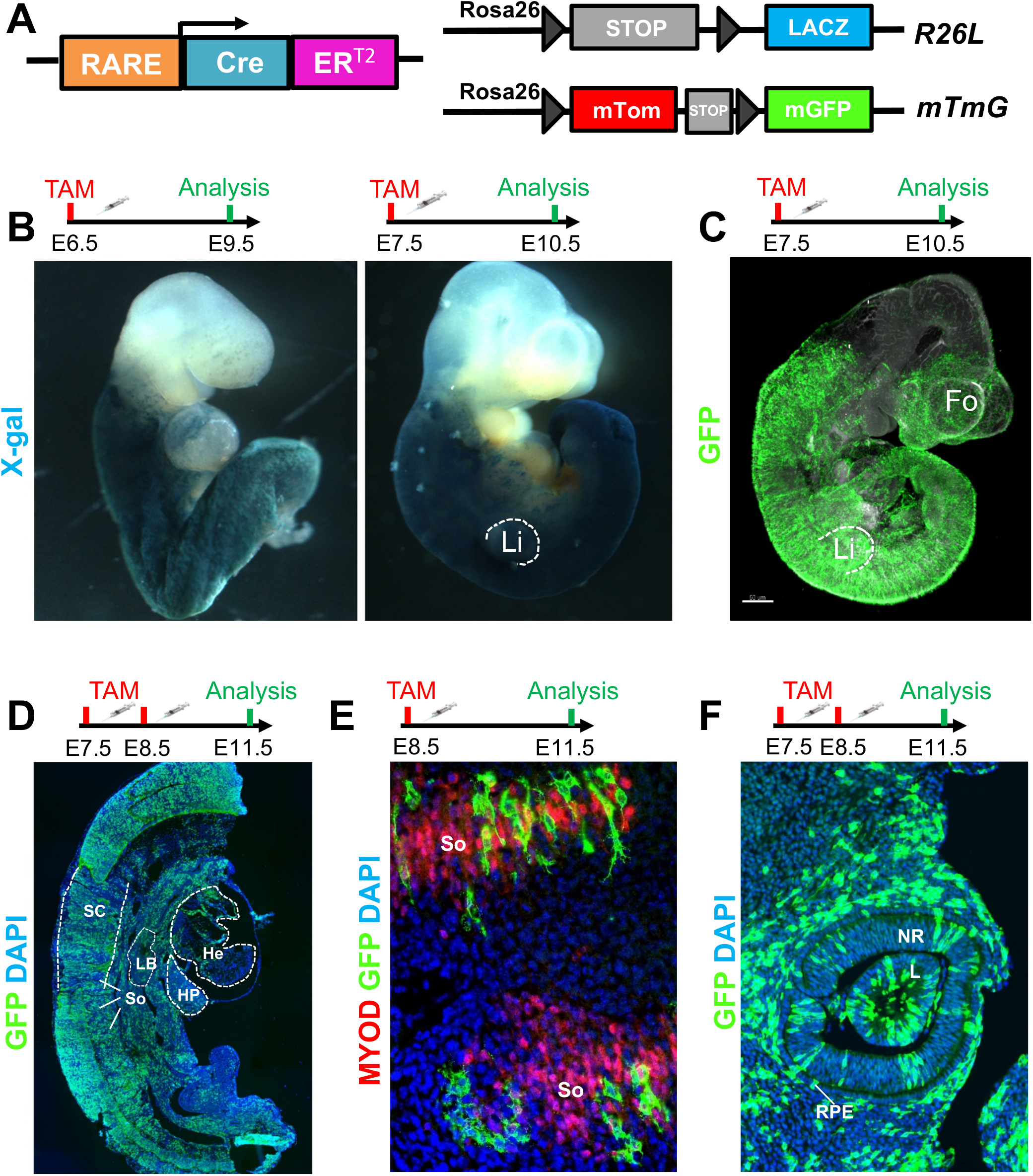
A novel *RARECreER*^*T2*^ line recapitulates endogenous RA signaling in mouse embryos. **(A)** Scheme illustrating the strategy used to test the novel *RARECreER*^*T2*^ line during various stages of embryonic development. The *RARECreER*^*T2*^ line was crossed with the *Rosa26LacZ* (*R26L*) or the membrane targeted tandem dimer Tomato membrane targeted green fluorescent protein (*mTmG*) reporter lines and recombination was induced via tamoxifen (TAM) administration at multiple time-points. Grey arrowheads represent *loxP* sites. **(B)** Whole-mount X-gal staining of *RARECreER*^*T2*^ *R26L* embryos induced at embryonic day 6.5 (E6.5) or E7.5 followed by respective analyses at E9.5 or E10.5 demonstrates efficient labeling with the *lacZ* reporter. Notice the labeling of the forelimbs (Li) with the E7.5 pulse. **(C)** Whole-mount GFP IF of *RARECreER*^*T2*^ embryos crossed with the *mTmG* reporter, pulsed with tamoxifen at E7.5 and sacrificed at E10.5 reveals specific forebrain (Fo) labeling. **(D)** GFP IF on sagittal sections of E11.5 embryos pulsed with tamoxifen at E7.5 and E8.5. Labelling is detected in various organs and tissues with well described RA activity such as the heart (He), spinal cord (SC), hepatic primordium (HP), lung bud (LB) and developing somites (So). Heads were removed for analysis (n=3 embryos analyzed, representative embryo shown). **(E)** Co-IF with GFP and Myoblast determination protein (MYOD) antibodies reveals RA-responsive cells in developing somites when pulsed at E8.5 and analyzed at E11.5. **(F)** Embryos pulsed with tamoxifen at E7.5 and E8.5 and analyzed at E11.5 display efficient labelling of the neural retina (NR), lens (L) and retinal pigment epithelium (RPE) of the eye as determined by GFP IF.

Another organ with a very well characterized RA response is the eye. *Aldh1a2* is detected in the optic vesicle as early as E8.5, where it is required for the transition from optic vesicle to optic cup (*45*-*47*). As development proceeds, *Aldh1a1* expression in the dorsal retina and *Aldh1a3* expression in the surface ectoderm, lens placode, retinal pigment epithelium and, later, the ventral retina, continue to activate RA signaling in the eye (*26*, *28*, *48*, *49*). Analysis of *RARECreER*^*T2*^ embryos pulsed with tamoxifen at E7.5 and E8.5 and analyzed at E11.5 revealed efficient GFP labeling of the lens placode, neural retina and prospective retinal pigment epithelium cells, consistent with previously published data (Fig. 1F) (*47*). Embryos pulsed at E8.5 and analyzed at E11.5 also displayed labeling in the eye, although at a lower frequency. These results are consistent with the fact that RA activity in the eye is continuous during initial stages of its development, since multiple pulses of tamoxifen lead to more efficient labelling when compared to only one shot of tamoxifen (Fig. S1A) Analysis of embryos pulsed at E8.5 and analyzed either at E10.5 (*R26L*) or E11.5 (*mTmG*) revealed labeling in the eyes, forebrain, heart and somites, but the overall labelling efficiency was decreased when compared to earlier time-points (Fig. S1B; S1C). Tamoxifen administration at E10.5 followed by light sheet microscopy analysis of whole embryos at E12.5 revealed a similar GFP labelling pattern as the E8.5 pulses (Fig. S1D). In summary, the *RARECreER*^*T2*^ line efficiently labels organs and tissues with well described RA activity in a highly dynamic fashion, and its labelling efficiency peaks during early stages of embryogenesis.

### Cardiomyocytes are highly responsive to RA signaling during embryonic development

RA signaling is essential for cardiac formation and displays unique spatiotemporal patterns of activity in the heart (*50*, *10*, *11*). During early stages of cardiac development, RA signaling is active in the venous pole of the heart. At later time-points, RA activity relocates to the heart’s outer layer, otherwise known as the epicardium (*51*, *52*). A closer look at the hearts of E9.5 *RARECreER*^*T2*^ embryos stained with X-gal after TAM treatment at E6.5 revealed specific labeling of the venous pole derivatives (atria and outflow tract) (Fig. 2A; S2A), and minimal labelling of the developing ventricles, faithfully recapitulating the pattern observed with other RA reporters (*5*, *6*). Meanwhile, embryos pulsed with tamoxifen at E10.5 and analyzed at E13.5 exhibited very strong labeling of the heart ventricles with almost no labeling of the atria and outflow tract (Fig. 2B), once again consistent with previously published data (*51*). Interestingly, analysis of E13.5 heart sections counterstained with eosin revealed the majority of the X-gal labeling to be within the myocardial wall of the heart and not the epicardium (Fig. S2B).

**Fig. 2:**
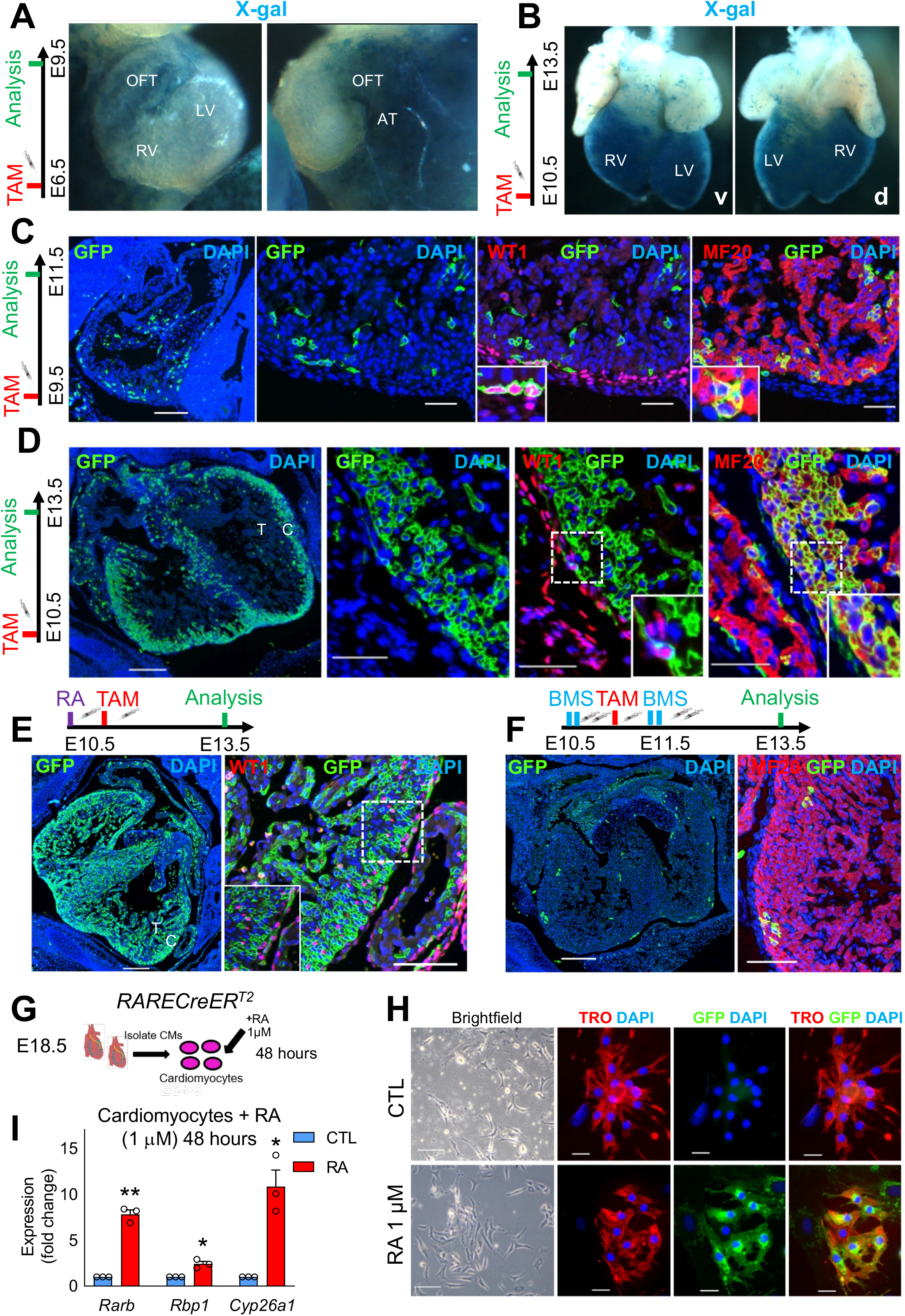
Cardiomyocytes are highly responsive to RA signaling during embryonic development. **(A)** Whole-mount X-gal staining of *RARECreER*^*T2*^ embryos pulsed with tamoxifen at E6.5 and sacrificed at E9.5 reveals strong labeling of venous pole derivatives of the heart (outflow tract (OFT) and atria (AT)). Minimal labeling is detected in the right ventricle (RV) and left ventricle (LV) of hearts. **(B)** Whole-mount X-gal staining of embryos pulsed with tamoxifen at E10.5 and sacrificed at E13.5 reveals strong ventral (v) and dorsal (d) labeling of the heart ventricles with minimal labeling of the atria and outflow tract. **(C)** Administration of tamoxifen (TAM) at E9.5 to *RARECreER*^*T2*^; *mTmG* embryos followed by analysis at E11.5 reveals specific labeling of the epicardium (GFP^+^WT1^+^ cells) and myocardium (GFP^+^MF20^+^ cells) in developing hearts. **(D)** Tamoxifen administration at E10.5 followed by analysis at E13.5 reveals strong labeling of the compact myocardium (C) and minor labelling of the trabecular layer (T) in developing hearts. Minimal labeling of the epicardium (WT1^+^ cells, inset) is detected at this time-point. Insets shown are from right ventricle of representative heart. **(E)** Exogenous supplementation of all-trans Retinoic acid (RA) (10 mg/kg) to pregnant dams 4 hours prior to tamoxifen induction leads to increased labeling of the trabecular myocardium and epicardium (WT1^+^ cells, inset) in developing hearts at E13.5 when compared to non-RA-treated embryos in **(D)** (n=3 embryos analyzed). **(F)** Supplementation of the RAR reverse agonist BMS493 (5 mg/kg) to pregnant dams 4 hours before and 4 hours after tamoxifen induction (extra two doses given in 8 hour interval 1 day after TAM induction) drastically reduces the number of GFP^+^ cells in *RARECreER*^*T2*^ hearts (n=3 embryos analyzed). **(G)** Schematic illustrating strategy for isolating primary cardiomyocytes (CMs) from hearts of E18.5 *RARECreER*^*T2*^; *mTmG* embryos followed by 48 hour treatment with 1 μM RA. **(H)** *RARECreER*^*T2*^; *mTmG* primary cardiomyocytes respond directly to RA treatment as demonstrated by co-IF for GFP and Troponin T. No GFP staining is detected in DMSO-treated control (CTL) cells. **(I)** qPCR analysis of primary cardiomyocytes isolated from E18.5 hearts and treated with 1μM RA leads to an upregulation of RA transcriptional targets (*Rarb, Rbp1, Cyp2a1*). Data are expressed as fold change vs. controls and columns are means ± SEM (n=3). WT1 = Wilms’ tumour protein, TRO = Troponin T, MF20 = Myosin heavy chain. Scale bars mosaics: 100 μM, Close ups: 40 μM. All statistics two tailed t-test assuming unequal variance, *p<0.05, **p<0.01.

To more precisely determine the cell-types labelled by the *RARECreER*^*T2*^ line during mid stages of gestation, we conducted lineage tracing experiments with the *mTmG* reporter allele, and then performed co-IF with GFP and epicardial (WT1 (Wilms’ tumour protein)) or myocardial (MF20 (myosin heavy chain))-specific antibodies. Administration of tamoxifen at E9.5 followed by analysis at E11.5 revealed GFP localization in both WT1^+^ and MF20^+^ cells (Fig. 2C). Meanwhile, tamoxifen administration at E10.5 followed by analysis at E13.5 revealed high labeling of MF20^+^ cardiomyocytes, but minimal labeling of WT1^+^ epicardial cells (Fig. 2D). The cardiomyocyte-specific labeling was highest in the compact layer, with minimal GFP^+^ cells observed in trabecular cardiomyocytes (Figure 2D, insets). Taken together, these data demonstrate that as heart development proceeds, cardiomyocyte precursors of the compact layer increasingly become the major RA-responsive cell-type.

To validate that the activation of GFP in cardiomyocytes from our *RARECreER*^*T2*^ embryos reflects true RA signaling, we performed various *in vivo* and *in vitro* experiments. Administration of RA to pregnant dams at E10.5 four hours prior to TAM induction increased the RA-response in *RARECreER*^*T2*^ hearts analyzed at E13.5, even labeling a substantial portion of trabecular myocytes (Fig. 2E). Conversely, treatment of embryos with the RAR inverse agonist BMS493 (*53*) prior to and following tamoxifen administration led to a very strong decrease in GFP^+^ cells (Fig. 2F). Moreover, isolation of primary cardiomyocytes from E18.5 *RARECreER*^*T2*^ hearts followed by RA treatment led to specific labelling of Troponin T^+^ cardiomyocytes (Fig. 2G, schematic; 2H). The efficiency of the RA treatment in cultured cardiomyocytes was confirmed by qPCR analysis, which showed upregulation of the RA target genes *Rarb*, *Rbp1* (retinol binding protein) and *Cyp26a1* (cytochrome P450 family 26 subfamily a polypeptide 1) ((Fig. 2I).

To ensure the cardiomyocyte response was not an artefact of our transgenic line, we performed lineage tracing experiments with a second *RARECreER*^*T2*^ line (Line B). Administration of tamoxifen at E10.5 followed by analysis at E14.5 revealed a nearly identical pattern of GFP staining in the compact myocardium when compared to our original line (Fig. S2C). Altogether, these data demonstrate that the cardiomyocyte-specific labeling observed with the *RARECreER*^*T2*^ line is a reliable response that can be influenced by exogenous activation or inactivation of the RA pathway.

### Cardiomyocyte-specific RA signaling is active during late stages of heart development

Although the role of RA signaling during heart looping and myocardial compaction is well described, it is unclear whether retinoids continue to play an important role after E13.5 (*9*). We next decided to look at later time-points of heart development by administering tamoxifen at E14.5 and sacrificing embryos at E18.5. Strikingly, we observed a high number of GFP^+^ cells, mainly cardiomyocytes (Fig. 3A). We also noticed that many of the labelled cells were located deep within the compact layer (Fig. 3A, white arrows), a striking observation given that at these time-points ALDH1A2 protein is believed to be restricted to the epicardium (*51*). To see if this deep labeling was simply due to clonal expansion of cardiomyocytes located near the epicardium, we traced cells for a shorter period of time by giving a pulse at E15.5 and analyzing at E17.5. Once again, we observed cardiomyocyte-specific labeling, both near the epicardium as well as deeper within the myocardial wall (Fig. 3B). Many of the GFP^+^ cardiomyocytes located away from the epicardium were separated from other cells (Fig. 3B, white arrows), suggesting their response arises from a local source of RA, independently of the epicardium.

**Fig. 3:**
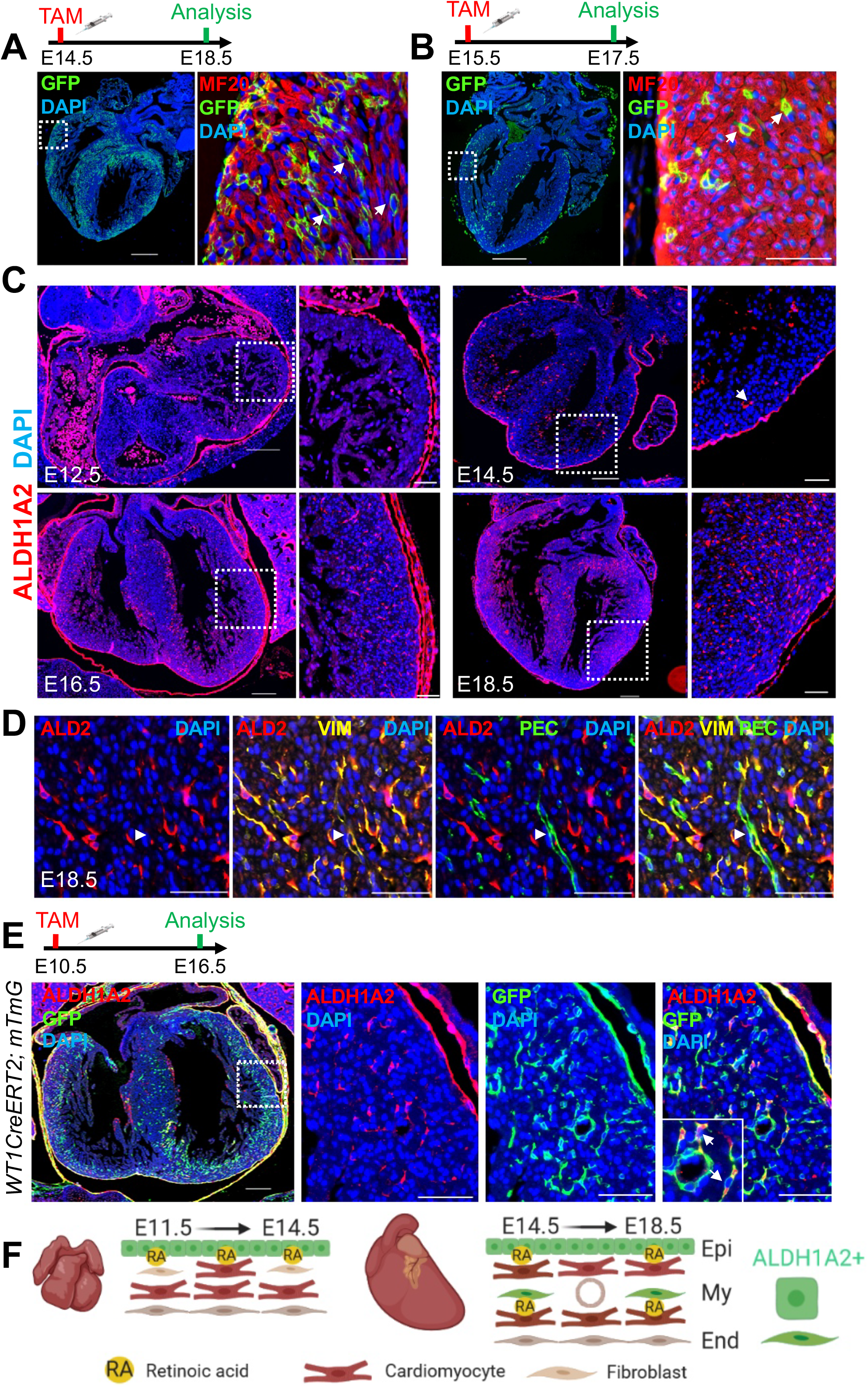
Cardiomyocyte-specific RA signaling is active during late stages of heart development. **(A)** Administration of tamoxifen (TAM) to *RARECreERT; mTmG* embryos at E14.5 followed by analysis at E18.5 reveals cardiomyocyte-specific (MF20^+^) labeling in developing hearts. Cardiomyocytes located deep within the ventricular wall are also labelled (white arrows). **(B)** Tamoxifen administration at E15.5 labels cardiomyocytes deep within the ventricular wall (white arrows) when analyzed only two days later at E17.5. **(C)** IF analysis reveals ALDH1A2 protein is restricted to the epicardium of the developing heart at E12.5. At E14.5 ALDH1A2 protein is detected within the ventricular wall (white arrow). High ALDH1A2 protein levels are detected in the ventricular wall at E16.5 and E18.5 (at least 3 embryos analyzed per time-point). **(D)** Co-IF with Vimentin (VIM) and PECAM antibodies reveals ALDH1A2 (ALD2) is produced by cardiac fibroblasts/connective tissue (VIM+) and not by endothelial cells (VIM+PECAM+, white arrowheads) in the developing heart. Images taken from representative region of interventricular septum. **(E)** Administration of tamoxifen to embryos carrying the *WT1CreER*^*T2*^ (epicardial-specific *CreER*^*T2*^ line) and *mTmG* alleles followed by analysis at E16.5 reveals many of the ALDH1A2^+^ cells within the ventricular wall of the developing heart are derived from the epicardium (GFP^+^). **(F)** Scheme illustrating pattern of ALDH1A2 production and RA activity in the myocardium during mid-late stages of cardiac development. Image created with Biorender software. Epi = epicardium, My = myocardium, End = endocardium. Scale bars mosaics: 100 μM, Close ups: 40 μM.

During cardiac formation, ALDH1A2 is the main enzyme responsible for RA synthesis and its loss of function leads to cardiac defects at both early and mid-stages of development (*40*, *14*, *54*). Whether or not ALDH1A2 has a role at later time-points is unknown since full *Aldh1a2*-null and *Aldh1a2*-null embryos rescued by a short-term RA supplementation die at E9.5 and E13.5, respectively (*40, 14*). To see if the RA response detected after E13.5 was due to *Aldh1a2* expression, we performed IF with an anti-ALDH1A2 antibody at various stages of gestation. At E12.5 ALDH1A2 was, as previously shown, restricted to the epicardium. After E14.5, however, ALDH1A2 could be detected within the ventricular and interventricular walls. This expression pattern increased at later time-points, peaking at E18.5 (Fig. 3C). To determine which cell-types were positive for ALDH1A2 we performed co-IF with various antibodies. No co-IF of ALDH1A2 with cardiomyocyte (MF20) (data not shown), smooth muscle (Transgelin) (data not shown) or endothelial (PECAM1) (Fig. 3D) markers was detected. By contrast, ALDH1A2^+^ cells were positive for the intermediate filament marker Vimentin, suggesting them to be cardiac fibroblasts or connective tissue (Fig. 3D). We next wanted to see if the ALDH1A2^+^ cells arose from the epicardium, which would be consistent with them being fibroblasts or other connective tissue cells. To this end, we performed lineage tracing experiments with the *Wt1*^*tm2(cre/ERT2)Wtp*^ line (referred to as *WT1CreER*^*T2*^) (*55*), which specifically labels the epicardium, crossed with the *mTmG* reporter. Tamoxifen induction at E10.5 followed by analysis at E16.5 revealed a significant portion of ALDH1A2^+^ cells within the ventricular wall were labelled with GFP, demonstrating they indeed originate from the epicardium (Fig. 3E, white arrows). Taken together, these data suggest that epicardial cells and epicardial-derived cardiac fibroblasts that have migrated into deeper layers of the forming myocardium produce ALDH1A2, and that the local RA signal is received by neighboring cardiomyocytes (Fig. 3F).

### The *RARECreER*^*T2*^ line labels several cell-types, including cardiomyocytes, in adult hearts subjected to myocardial infarction

RA signaling appears to play a protective role in the adult heart after acute damage (*20*, *21*, *56*). To further analyze RA signaling in cardiac repair, we subjected *RARECreER*^*T2*^ mice to surgical ligation of the left coronary artery (a widely used experimental myocardial infarction model). To label RA-responsive cells, one pulse of tamoxifen was given immediately after surgery, and a second pulse 48 hours later (Fig. 4A, schematic). Strikingly, analysis of hearts six days after MI revealed drastic GFP enrichment in infarct hearts when compared to sham controls (Fig. 4B). The staining pattern of GFP closely overlapped with ALDH1A2, suggesting it to be the major enzyme driving the RA response in infarct hearts, which is consistent with previous studies (*25*). *Aldh1a1* and *Aldh1a3* were also upregulated as determined by qPCR analysis (Fig. 4C). A closer look at the staining patterns revealed the ALDH1A2^+^ and GFP^+^ cells increased particularly within the injury and border zones of damage (Fig. 4B; 4F). Most ALDH1A2^+^ cells were GFP-negative, suggesting a paracrine rather than autocrine mode of action (Fig. 4D). Co-IF for GFP with various markers demonstrated that many different cell-types were responsive to RA. These included PECAM1^+^ coronary vessels, αSMA^+^ activated fibroblasts and/or smooth muscle cells and Troponin T^+^ cardiomyocytes (Fig. 4E). Interestingly, a substantial portion of cardiomyocytes in the injury border zone, a region that is highly remodelled after MI, were also GFP^+^ (Fig. 4F, white arrows).

**Fig. 4:**
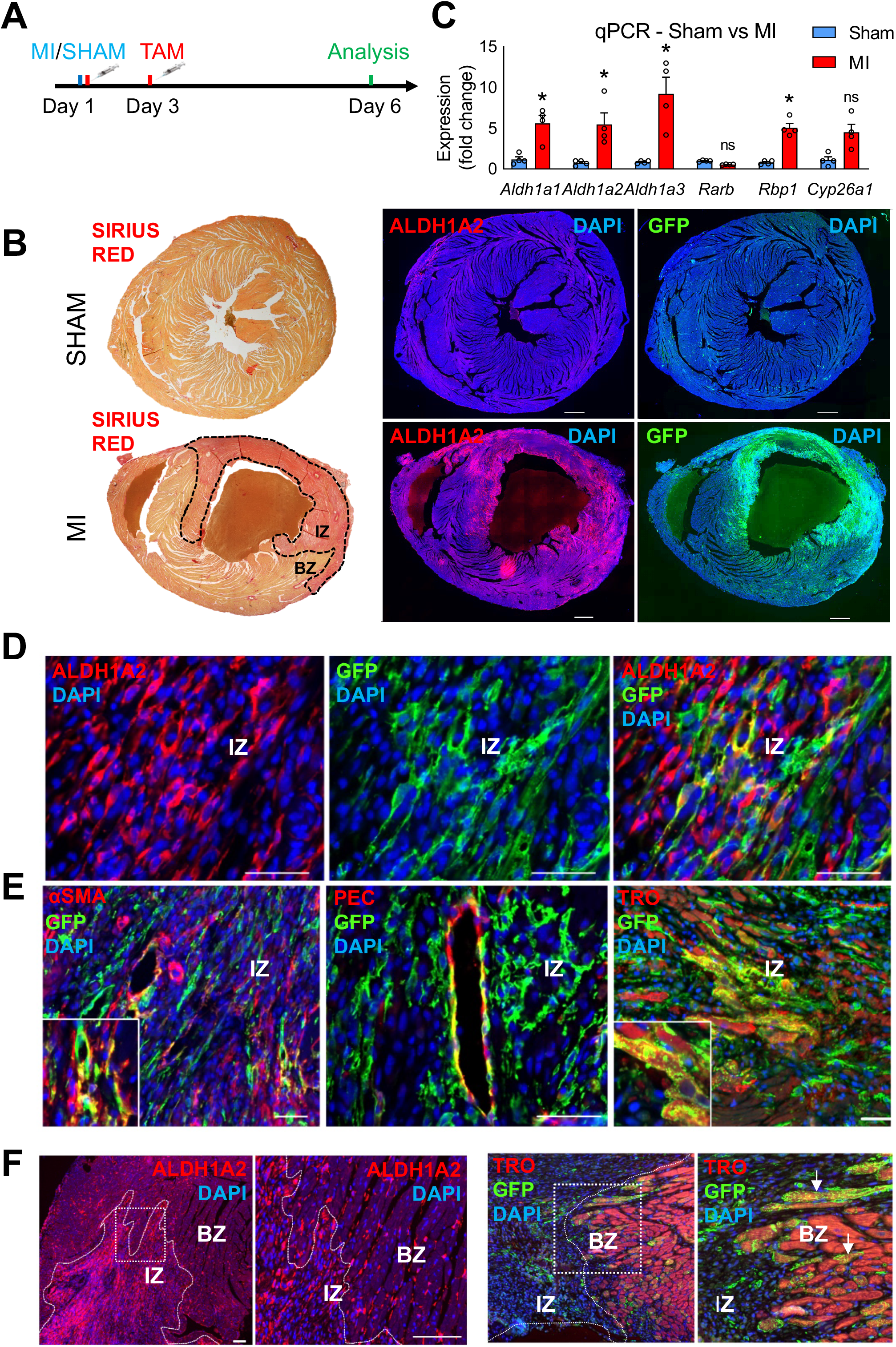
The *RARECreER*^*T2*^ line labels several cell-types, including cardiomyocytes, in adult hearts subjected to myocardial infarction. **(A)** Schematic illustrating the lineage tracing experiments performed in *RARECreER*^*T2*^; *mTmG* mice subjected to myocardial infarction (MI). Tamoxifen was administered twice (30 minutes and 48 hours after surgery) and hearts were analyzed 6 days post MI. **(B)** IF on infarct and sham hearts from *RARECreER*^*T2*^ mice reveals ALDH1A2 and GFP are highly enriched in and around the infarct zone (IZ) (marked by Sirius Red staining, dotted black line) while minimal ALDH1A2 and GFP staining is detected in sham hearts. BZ = border zone of injury. **(C)** qPCR analysis on RNA extracted from infarct and sham hearts reveals *Aldh1a1,2* and *3* and *Rbp1* are upregulated after MI. The RA targets *Rarb* and *Cyp26a1* are not significantly altered. Data are expressed as fold change vs controls and columns are means ± SEM (n=4 hearts). **(D)** Co-IF for ALDH1A2 and GFP demonstrates minimal co-staining in *RARECreER*^*T2*^ MI hearts. **(E)** Co-IF for GFP plus αSMA, PECAM1 or Troponin T demonstrates an RA response in activated fibroblasts/smooth muscle cells, coronary vessels and cardiomyocytes respectively in *RARECreER*^*T2*^ MI hearts. **(F)** Closer analysis of *RARECreER*^*T2*^ infarct hearts reveals ALDH1A2 protein localization to the infarct zone and border zone. GFP^+^ cells also localize to both regions, and many of the GFP cells in the border zone are cardiomyocytes as demonstrated by co-IF for Troponin T (white arrows). TRO = Troponin T, SMA=smooth muscle actin. All statistics two tailed t-test assuming unequal variance, *p<0.05, ns = not significant. Scale bars: mosaics 100 μM, close ups 40 μM.

### Depletion of RA signaling leads to larger infarct zones and increased apoptosis

Since our *RARECreER*^*T2*^ line showed a strong response in regions of the heart that are highly remodelled during cardiac repair, we next wanted to study the effects of MI on mice with reduced levels of RA signaling. For this we used mice carrying both floxed *Aldh1a1/a2/a3* alleles and the *CAGGCreER*^*TM*^ transgene. To delete the enzymes, we injected mice five times with tamoxifen one week prior to surgery (Fig. 5A, schematic). Analysis of hearts six days after MI revealed reasonable deletion efficiency, with a 50% reduction in ALDH1A2 protein (Fig. 5C) and a near 70% decrease of *Aldh1a1* and *Aldh1a2* mRNA levels (Fig. 5D). Incomplete deletion of *Aldh1a* genes was likely due to the inaccessibility of their loci, as testing of the *CAGGCreER*^*TM*^ crossed with the *mTmG* line showed very efficient recombination in the adult heart (Fig. S3). The resulting mutant mice were nonetheless referred to as *Aldh1a*-knockout mice (*RAKO*).

**Fig. 5:**
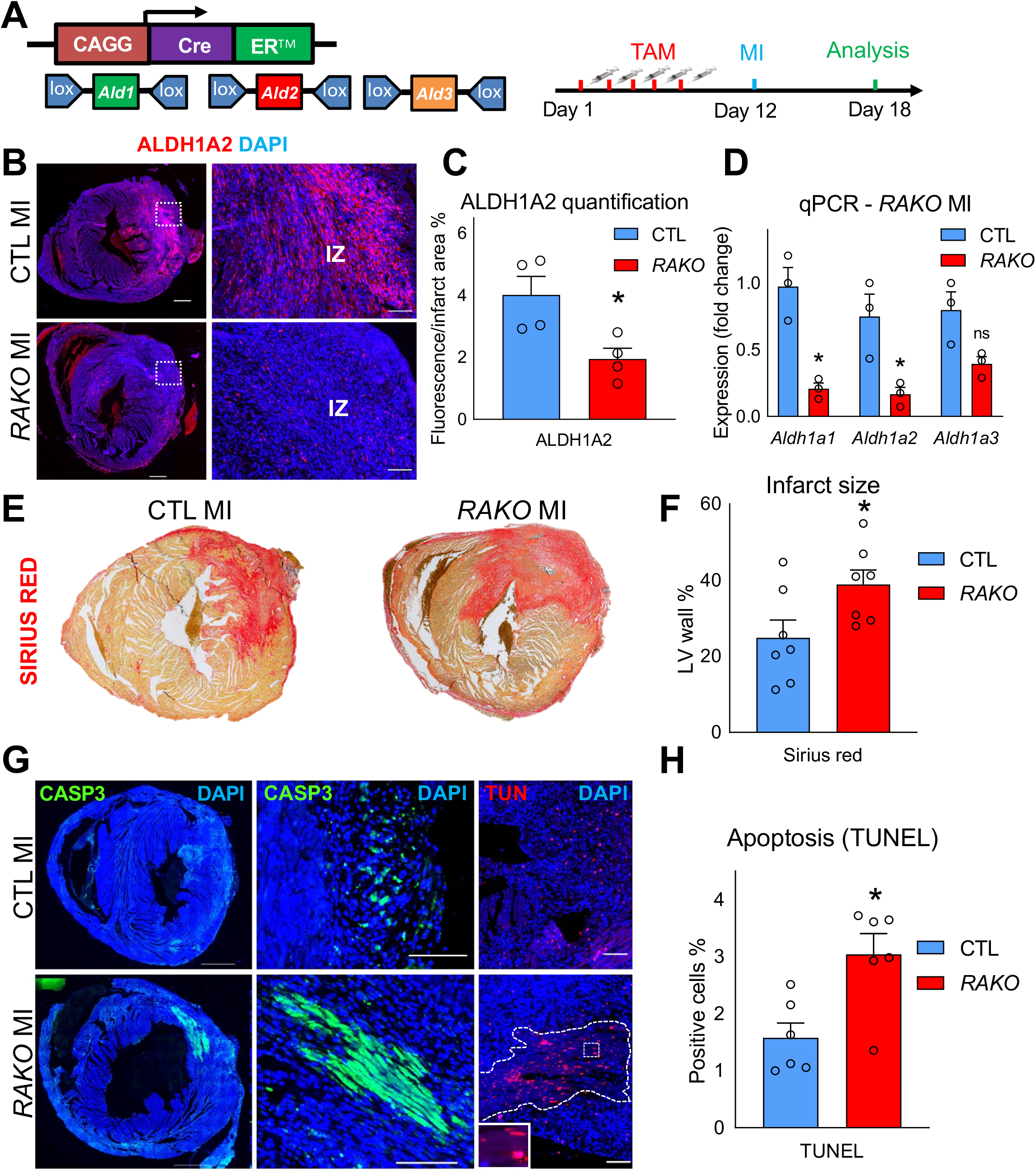
Depletion of RA signaling leads to larger infarct zones and increased apoptosis. **(A)** Schematic illustrating strategy used to delete floxed alleles of the *Aldh1a1,2,3 (Ald1,2,3)* enzymes with the *CAGGCreER*^*TM*^ line (mutant mice referred to as *RAKO*s). Five daily doses of tamoxifen were administered one week prior to surgery and operated hearts were analyzed 6 days post MI. **(B)** IF analysis reveals a significant decrease of ALDH1A2 protein in *RAKO*s when compared to *CAGGCreER*^*TM*^ negative (control (CTL)) hearts. **(C)** Quantification of ALDH1A2 protein levels by IF reveals a 50% decrease in *RAKO* hearts. ALDH1A2 pixel area was divided by total infarct area and measurements were performed using ImageJ. Columns are means ± SEM (n=4 hearts). **(D)** qPCR analysis of RNA extracted from infarct hearts reveals significant decreases in *Aldh1a1* and *Aldh1a2* expression in *RAKO*s when compared to controls. *Aldh1a3* expression is also reduced, though not significantly. Data are expressed as fold change vs. controls and columns are means ± SEM (n=3 hearts). **(E)** Sirius red detection of collagen deposition demonstrates increased infarct size in *RAKO* hearts when compared to controls. **(F)** Quantification of infarct size in *RAKO* and control hearts. The infarct areas were measured with ImageJ software and were normalized to the total area of the left ventricle. Columns are means ± SEM (n=7 hearts). **(G)** Active caspase 3 and TUNEL stainings reveal increased apoptosis in *RAKO* hearts when compared to controls. *RAKO* hearts have visible “patches” of apoptotic cells (lower middle panel; white outline in lower right panel). **(H)** Quantification of TUNEL^+^ cells in *RAKO* and control infarct hearts using ImageJ software. Columns are means ± SEM (n=6 hearts). CASP3 = active caspase 3, TUN = TUNEL, IZ = infarct zone. All statistics two tailed t-test assuming unequal variance, *p<0.05, ns = not significant. Scale bars: mosaics 100 μM, close ups 40 μM.

To analyze the effects of depleting the RA pathway prior to MI wse measured the size of the infarct zones in *RAKO* and control mice. Strikingly, *RAKO* mice exhibited significantly increased infarct zones as revealed by collagen staining with Sirius red (Fig. 5E; 5F). Furthermore, *RAKO* mice showed increased rates of apoptosis as determined by active caspase 3 and TUNEL (Terminal deoxynucleotidyl transferase dUTP nick end labeling) staining (Fig. 5G; 5H). No apoptosis was observed in RAKO mice subjected to sham surgery (Fig. S4A). Interestingly, many of the apoptotic cells in *RAKO* mice were cardiomyocytes and they tended to concentrate in clusters or so called “patches” that were positive for both active caspase3 and TUNEL (Fig. 5G; S4C). These patches were observed in four out of seven *RAKO* mice and only one out of eight control mice (Fig. S4D, table). Thus, dampening the RA response prior to MI leads to increased cardiomyocyte apoptosis, suggesting RA signaling plays a protective role in damaged heart muscle.

A previous study using exogenous RA supplementation in mice subjected to ischaemia/reperfusion determined that the RA pathway reduced apoptosis in damaged hearts by modulating the MAP kinase pathway. This was suggested to be through direct transcriptional regulation of *Adam10*, a gene which encodes for a protease that cleaves and inactivates the RAGE10 (Receptor for Advanced Glycation End products) receptor, a positive regulator of the ERK1/2 MAP kinase pathway (*21*). However, we did not observe any differences in ERK1/2 activation, as determined by IF for phospho-ERK1/2, in our *RAKO* mice (Fig. S5A; S5B). Additionally, treatment of primary cardiomyocytes with RA or BMS493 did not lead to significant changes in *Adam10* expression (Fig. S5C; S5D). Taken together, these data suggest the anti-apoptotic effects of RA observed in our model of RA depletion to be unrelated to *Adam10*/MAP-Kinase signaling.

### RA treatment in embryonic cardiomyocytes promotes a notable transcriptional response and regulates genes involved in cardiac repair, including *Tgm2* and *Ace1*

The labeling of cardiomyocytes with our *RARECreER*^*T2*^ line, as well as the increase in apoptosis observed in *Aldh1a-*null mice after MI led us to hypothesize that RA signaling plays a protective role specifically in cardiomyocytes. To decipher the underlying mechanisms, we isolated primary cardiomyocytes from E18.5 hearts and treated them with 100 nM RA for 48 hours to mimic long-term exposure to RA, which is normally experienced after MI. We then extracted RNA and used the samples for high-throughput sequencing (Fig. 6A, schematic). Analysis of the sequencing data identified several canonical RA targets such as *Rarb*, *Cyp26b1* and *Cyp26a1* to be top hits among all genes analyzed (Fig. 6B, Table S1), and gene ontology analysis revealed enrichment of GO terms for inflammatory and DNA damage responses (Fig. 6C). Furthermore, a handful of genes previously shown to be important in cardiac development and MI were also determined to be either up or downregulated (Fig. 6B). Of note, *Tgm2*, which has been shown to promote ATP synthesis and thus limit damage after MI, was significantly upregulated (Fig. 6B) (*57*). We also noticed that RA treatment significantly repressed the expression of Angiotensin converting enzyme 1 (*Ace1*). ACE1 is responsible for converting angiotensin 1 into angiotensin 2, which in turn activates the renin angiotensin system and subsequent vasoconstriction. Upregulation of ACE1 has been observed in rodent hearts after MI and is generally considered to be harmful (*58*). Indeed, ACE inhibitors are commonly used to treat patients recovering from MI, and they have shown great success in various clinical trials (*59*). Moreover, a connection between RA signaling, the RAS system, and cardiomyocyte-specific apoptosis has been previously established *in vitro* (*60*, *22*). Activation and repression of *Tgm2* and *Ace1*, respectively, was confirmed by qPCR analysis (Fig. 6D). To ensure the regulation of *Tgm2* and *Ace1* was not due to downstream indirect effects, we treated primary cardiomyocytes with RA for only nine hours and thereby observed significant modulation of both genes (Fig. 6E). We also treated cardiomyocytes with BMS493 for 48 hours, and observed that *Tgm2* expression was repressed while *Ace1* levels were increased, consistent with our hypothesis that both genes are regulated by RA signaling (Fig. 6F). *Ace1* is highly expressed by endothelial cells in the heart and it is possible the repression of *Ace1* by RA may be a result of contaminating endothelial cells. To address this issue, we removed endothelial cells from our cardiomyocyte cultures using CD31 magnetic beads, then treated enriched cardiomyocytes with RA (Fig. S5E; S5F). Consistently, RA treatment repressed *Ace1* expression (Fig. 6G). Taken together, these data show that RA can stimulate a significant transcriptional response in primary cardiomyocytes, and that some of the regulated genes, such as *Tgm2* and *Ace1*, are known to play important roles in cardiac repair.

**Fig. 6:**
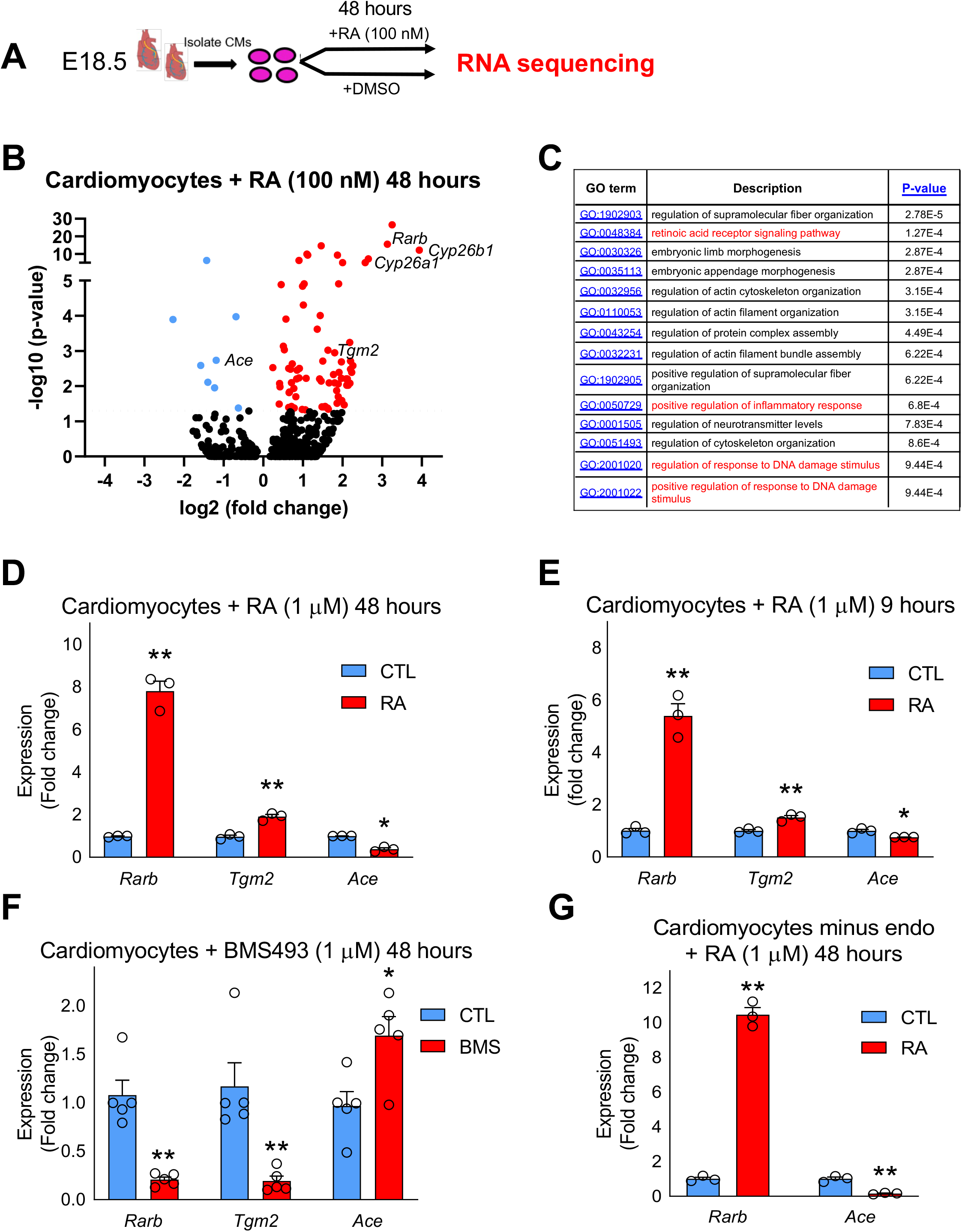
RA treatment in embryonic cardiomyocytes promotes a notable transcriptional response and regulates genes involved in cardiac repair such as Tgm2 and Ace1. **(A)** Schematic illustrating the strategy used to isolate primary cardiomyocytes from E18.5 hearts. Cultured cardiomyocytes were treated for 48 hours with 100 nM RA. RNA was extracted, libraries were prepared with oligo (dT) primers and single end sequencing was performed on RA-treated and DMSO-treated cells (n=4 biological replicates per treatment). **(B)** Volcano plot analysis of RNA sequencing results performed with Graphpad software. Dotted line represents significance threshold equivalent to p<0.05. Several canonical RA targets (*Rarb; Cyp26a1*; *Cyp26b1*; red) are significantly upregulated. Genes involved in cardiac repair such as Transglutaminase 2 (*Tgm2)* (red) and Angiotensin converting enzyme 1 (*Ace1*) (blue) are also significantly altered. DESeq analysis of genome aligned reads was performed using proprietary Genomatix software. Only the top 500 genes were included in the volcano plot analysis. **(C)** Gene ontology (GO) analysis of biological processes using publically available Gorilla software. Only the top 500 genes were included in the GO analysis. **(D)** qPCR analysis confirms upregulation of *Tgm2* and repression of *Ace1* mRNA levels in primary cardiomyoctes treated with 1 μM RA for 48 hours (n=3). **(E)** Acute nine hour 1 μM RA treatment in primary cardiomyocytes promotes *Tgm2* upregulation and *Ace1* repression as shown by qPCR analysis (n=3). **(F)** Treatment of primary cardiomyocytes with the RAR signaling reverse agonist BMS493 (1 μM) for 48 hours reveals a decrease in *Tgm2* expression and an increase in *Ace1* expression levels as shown by qPCR analysis (n=5). **(G)** Removal of endothelial cells (endo) with CD31-magnetic beads followed by 48 hour 1 μM RA treatment on purified primary cardiomyocytes reveals significant repression of *Ace1* expression as shown by qPCR analysis (n=3). For all graphs data are expressed as fold change vs. controls and columns are means ± SEM. *Tgm2* = Transglutaminase 2, *Ace* = angiotensin converting enzyme. All statistics two tailed t-test assuming unequal variance, *p<0.05, **p<0.01.

## DISCUSSION

Here we have characterized a novel, tamoxifen inducible reporter for RA signaling in mice that can be used to permanently label RA-responsive cells and trace their descendants in specific tissues and organs. By performing several inductions and analyzing embryos at different time-points during gestation, we show that our line faithfully recapitulates the pattern of RA activity observed with other well-characterized RA reporters. Notably, at embryonic stages, TAM-induced *RARE-CreER*^*T2*^ transgenics display specific labeling in the somites, limbs, forebrain, heart, and caudal regions of the body, consistent with well described roles for the RA pathway (*1*, *3*, *4*). We also characterize a novel and direct RA response in cardiomyocytes during multiple phases of embryonic development, as well as after MI.

An interesting finding of our study is the highly dynamic response of the *RARECreER*^*T2*^ line in various organs when activated at different time-points. Tamoxifen administration at E6.5 and E7.5 revealed very efficient labelling of multiple organs and tissues, while pulses given at later time-points such as E8.5 and E10.5 led to reduced recombination in most regions of the embryo. Overall, these data are consistent with previously described early roles of RA signaling in early embryonic germ layers and in body axis patterning, limb bud formation and neural tube patterning (*34*, *41*-*43*). Other organs that have continuous RA activity throughout development such as the retina, somites, forebrain and heart can be labelled using the *RARECreER*^*T2*^ line with later pulses of tamoxifen (*2*, *3*). The biological relevance of these results not only validate our *RARECreER*^*T2*^ line, they also highlight its sensitivity and vast potential as a reliable tool for studying RA signaling and the fate of RA responding cells *in vivo*.

Previous studies have shown that RA signaling is involved in myocardial compaction but that RA does not act directly on cardiomyocytes (*15*, *16*). Instead, RA signaling was believed to act in an autocrine or extra cardiac manner in the epicardium and liver/placenta, respectively (*18*, *19*). Using our novel *RARECreER*^*T2*^ line, we clearly demonstrate that cardiomyocytes do indeed respond to RA. Moreover, we have validated this response through *in vivo* and *in vitro* modulation of the RA pathway. However, it is still not clear whether this response plays an important role in myocardial compaction. Deletion of the RXRA receptor in the myocardium does not lead to an obvious heart phenotype (*15*), but perhaps other receptors compensate for this loss of function. It is also possible that RA signaling may have essential roles in fine-tuning the patterning and maturation of the compact myocardium. One way of addressing this would be to perform a simultaneous myocardial-specific deletion of the three RARs. The deletion would have to target all three alleles since the RARs display high functional redundancy (*61*). Such a study would be useful in further delineating the roles of RA signaling during myocardial compaction and would provide additional insight into how RA regulates the proliferation, differentiation and/or maturation of cardiomyocytes.

The cardiomyocyte response as well as the dynamic pattern of ALDH1A2 protein localization observed during late stages of heart development is intriguing. Why do cardiomyocytes continue to respond to RA signaling at these time-points? Could RA be promoting the differentiation and maturation of these cells? Do the RA-responsive cells represent unique subpopulations with specific roles in the heart? All of these scenarios are plausible and should be addressed in future studies. The fact that ALDH1A2 protein is produced by alternative cell-types other than the epicardium is logical, since during later time-points, when the heart’s compact myocardial layer is thickened, it is unlikely RA can diffuse easily to cardiomyocytes located in deeper portions of the ventricular wall. Through a series of co-IF and lineage tracing experiments we suggest that the ALDH1A2^+^ cells at these time-points are cardiac fibroblasts or other connective tissue cells derived from the epicardium. However, more detailed analyses such as FACS analysis with multiple markers or single cell RNA sequencing would have to be carried out in order to confirm these findings. It would also be interesting to investigate if these same cell-types also produce ALDH1A2 after MI.

It is well known that RA signaling plays a protective role in the heart after acute damage. Treatment of rats with RA reduces cardiac hypertrophy and remodeling after MI and deficiency of RA leads to larger infarct zones (*56*, *20*). In mice, RA treatment also leads to smaller infarct zones and reduced apoptosis after ischaemia reperfusion (*21*). Our observation of increased scarring and apoptosis after infarct in *RAKO* mice is consistent with these studies. However, other studies suggest that the protective effects of RA signaling is context dependent, and that short-term localized exposure to retinoids after MI may even be harmful (*62*). In healthy non-infarcted mice, deletion of the RARA receptor has detrimental effects, mainly due to increased reactive oxygen species and calcium mishandling, while long-term RA treatment leads to cardiac hypertrophy and overall negative effects (*23*, *63*). Overall, these contradictory findings suggest that the effects of RA signaling in the heart are highly dependent on the dose and length of exposure to RA, meaning that for RA signaling to be protective, it needs to be tightly regulated. Our *RARECreER*^*T2*^ line can facilitate the monitoring of the RA response post MI. It can also help determining which cells respond to RA, as well as their respective fates in the long-term. Mechanistically, it is still unclear how RA promotes or inhibits cardiac repair. Some studies suggest RA plays an antioxidant role, while others suggest it inhibits the normally detrimental MAP kinase pathway (*23, 21, 60*). However, we could not find evidence for either hypothesis in our model of RA deficiency, and further work is necessary to discover the mechanisms responsible for the increased apoptosis observed in cardiomyocytes of *RAKO* mice.

To our knowledge, this is the first study to conduct RNA sequencing on primary cardiomyocytes treated with RA. Despite traditionally being considered non-responsive cells, RA treatment led to a notable transcriptional response in cardiomyocytes. In many ways this validates our observations with the *RARECreER*^*T2*^ line, in which cardiomyocytes, especially at later time-points are highly responsive to RA. However, these results need to be interpreted with caution since the cardiomyocytes used for the RNA sequencing were isolated from E18.5 hearts, whose response to RA may not accurately reflect the transcriptional landscape during initial phases of myocardial compaction (E10.5-E14.5) or after MI, in which the myocardium experiences severe hypoxia and inflammation. Furthermore, RA signaling can regulate gene expression post-transcriptionally and it is not clear if the targets identified from the RNA sequencing analysis are direct or indirect targets. Further work will be needed to clarify this. Nevertheless, given the fact that most of the targets tested responded to pharmacological activation or inhibition of RA signaling, it is likely that the RNA sequencing analysis is biologically relevant, and may, in future studies, help elucidate the role of RA signaling in the heart during late stages of development and after MI.

Of the many genes identified from the RNA sequencing analysis, *Tgm2* attracted our attention since it has previously been associated with regulating ATP synthesis after MI. In fact, *Tgm2* knockout mice exhibit larger infarcts when compared to controls, much like our *RAKO* mice (*57*). More recently, however, *Tgm2* has been shown to promote cardiac fibrosis in MI hearts and pharmacological inhibition of TGM2 led to smaller infarcts (*64*). It is thereby possible that the effects of *Tgm2* on heart repair, as with RA signaling, are dependent on the level of activity. Numerous links between RA signaling and the RAS pathway have been previously shown (*22*, *60*, *65*-*69*). Nevertheless, it is not entirely clear how RA signaling regulates RAS, and no direct RA transcriptional targets from the RAS pathway have yet been identified. We also show that *Ace1* expression is repressed by RA in cardiomyocytes, an important finding given that *Ace1* upregulation after MI has negative effects on cardiac remodeling (*58*). In fact, ACE inhibitors have consistently led to improved patient outcomes in clinical trials (*59*). Furthermore, one study has even shown that ACE1 can be detected in cardiomyocytes from human infarct hearts (*70*). It is thus possible that combining RA, which specifically inhibits *Ace1* expression, with ACE inhibitors may have synergistic beneficial effects in protecting cardiomyocytes post-MI, an interesting prospect for future studies aimed at improving the survival of patients suffering from heart disease.

In summary, we have shown that cardiomyocytes are responsive to RA signaling and that depletion of the RA pathway leads to increased cardiomyocyte apoptosis after MI. These findings have important implications in the heart development and cardiac regeneration research fields.

## MATERIALS AND METHODS

### Experimental Design

The aim of the study was to investigate the role of endogenous Retinoic Acid (RA) signaling in the heart using a novel transgenic mouse RA-reporter (*RARECreER*^*T2*^) and an *in vivo* genetic approach of deletion of *Aldh1a1, Aldh1a2* and *Aldh1a3* genes (i.e. mice lacking all sources of RA *in vivo*, an approach validated in (*71*)) to challenge the dogma that cardiomyocytes, the principle cell type of the heart, are not responsive to RA signaling both during embryonic development and after MI. The number of samples was determined on the basis of experimental approach, availability, and feasibility required to obtain definitive results. The numbers of replicates are specified in the appropriated *Materials and Methods* sections. The researchers were not blinded during data collection or analysis.

### Mice

All animal work was conducted according to national and international guidelines and was approved by the local ethics committee (PEA-NCE/2013/88). The *R26L*, *mTmG*, *WT1CreER*^*T2*^, *Aldh1a1*^*fl*^, *Aldh1a2*^*fl*^, *Aldh1a3*^*fl*^, and *CAGGCre-ER*™ lines have been described previously (*31*, *37*, *55*, *26*-*29*). For embryonic *RARECreER*^*T2*^ lineage tracing Cre activation was obtained by administration (gavage) of 200 mg/kg tamoxifen (Sigma-Aldrich) dissolved in corn oil (Sigma-Aldrich) to pregnant females at the indicated time-points. For adult myocardial infarction experiments (*RARECreER*^*T2*^ lineage tracing and *Aldh1a1/a2/a3* deletion experiments) Cre activation was obtained by administration (intraperitoneal injection) of 100 mg/kg tamoxifen at the indicated time-points. Details of RA and BMS493 *in vivo* treatments are provided in the figure legends.

For generation of *RARECreER*^*T2*^ mice, fertilized zygotes were obtained from super-ovulated B6D2F1 females mated to B6D2F1 males. Linearized DNA consisting of three copies of a 34 bp oligo (5) containing the direct repeat 5 (DR5) RARE of the murine *Rarb2* promoter upstream of the hsp68 promoter and *CreER*^*T2*^ gene was injected into the pronuclei of zygotes. Zygotes were then transferred into the oviducts of pseudo pregnant mice. All mice used in these experiments were heterozygous for the transgene, and were back-crossed for more than five generations. Two founder lines were established yielding the expected RA-responses during embryonic development, and the line with the higher activity was used for further study.

### Myocardial Infarction surgeries

Transgenic males 8 to 10-week-old were subjected to permanent ligation of left-anterior descending coronary artery. One dose of buprenorphine (0.1 mg/kg) was administrated subcutaneously for pre-operative analgesia. After 30 minutes they were anesthetized by inhalation of 4% isoflurane in a filled chamber and immediately intubated for artificial ventilation to maintain a respiratory rate of 135/min with pure oxygen mixed with 1-2% isoflurane. A left-sided thoracotomy was performed between the third and fourth ribs: the pericardium was cut open and ligation of the descending branch of the left coronary artery was made 2.5 mm under the tip of the left auricle using 8-0 silk suture. Sham operated mice underwent exactly the same procedure, except no ligation around the left coronary artery was made. Subsequently, the intercostal space, muscles of the external thoracic wall and skin were sutured with 6/0 polyester. All animals received 0.5 ml saline IP post-surgery to compensate for fluid loss and mg/kg of buprenorphrine subcutaneously for analgesia.

### Whole-mount X-gal staining

For whole-mount X-gal analysis, embryos or dissected organs were fixed for 45 minutes in 0.1% glutaraldehyde diluted in PBS, washed three times with wash buffer (2 mM MgCl2, 0.2% NP-40, 0.1% sodium deoxycholate in sodium phosphate buffer) and then incubated overnight (O/N) at 37°C in X-gal staining solution (washing buffer + 5 mM potassium ferrocyanide, 5 mM potassium ferricyanide and 1 mg/ml X-Gal substrate). For analysis on sections, samples were fixed O/N in 4% paraformaldehyde, embedded in paraffin, cut at 5 μM on a microtome, de-paraffinized and counterstained with eosin.

### Whole-mount GFP staining

Embryos were collected on their respective days and fixed with 4% paraformaldehyde overnight at 4⁰C. Embryos were dehydrated using an ascending concentration of methanol prepared in PBS, and bleached with 6% hydrogen peroxide O/N at 4⁰C. On the following day embryos were rehydrated using a descending concentration of methanol. Embryos were quenched with 0.3 M Glycine with Triton X (prepared in PBS) for 4-6 hours at room temperature. The Triton X concentrations were decided based on the stage of the embryos (0.5% Triton X concentration was used for E10.5 embryos and 1.0% Triton X concentration was used for E12.5 embryos). Samples were blocked in PBSGT (0.2% Gelatin (VWR), Triton X and 0.1 g/L of Thimerosal ((SIGMA), prepared in PBS) for 2 days at room temperature. Chicken Anti-GFP antibody and goat Anti-GATA4 were diluted in PBSGT at 1:500 concentrations. Samples were incubated with primary antibody solutions, 24 hours for E10.5 embryos and 1 week for E12.5 embryos at room temperature. Samples were washed six or more times over a day with PBSGT. Secondary antibodies were diluted in PBSGT at 1:500 concentration. Embryos were incubated in secondary antibodies for one (E10.5) to three (E12.5) day(s) at room temperature. Embryos were washed six or more times over a day with PBSGT.

### Image acquisition for whole-mount GFP staining

The protocol was adjected from a previously described protocol (*72*). Embryos were embedded in 1.0% low melting agarose (Invitrogen) prepared in PBS. The embedded embryos were transferred into amber colour glass vials and dehydrated with ascending concentrations of methanol. BABB solution was prepared by mixing one portion of benzyl alcohol (SIGMA) with two portions of benzyl benzoate (SIGMA). Samples were treated with 50% BABB diluted in methanol for 3-4 hours. Samples were then treated with the BABB solution until they were settled at the bottom of the vial. Samples were stored and mounted for imaging in BABB solution at room temperature. Images were acquired using Zeiss LSM780 microscopy (or homemade light-sheet microscopy) at the PRISM Imaging Facility. 3D constructions and analysis were performed using Imaris software.

### Immunofluorescence, histological analysis and Sirius red staining

For IF experiments, tissues were fixed overnight in 4% paraformaldehyde, progressively dehydrated and embedded in paraffin. 5 μM thick sections were rehydrated, boiled in a pressure cooker for 2 minutes with Antigen Unmasking Solution (Vector laboratories) and blocked in PBS solution containing 10% normal donkey serum and 3% BSA. All antibodies were applied overnight at 4°C at the concentrations listed in the antibody table (see Table S2). Secondary antibodies were diluted 1:400 and applied at room temperature for 1 hour. For histological analysis, 5 μM thick sections were stained with haematoxylin and eosin according to standard procedures. For Sirius red staining, 5 μM thick sections were stained in Sirius red solution (Sirius red powder (SIGMA) dissolved in picric acid) for 1 hour at room temperature. Samples were then washed in acidified water before being dehydrated with three 100% ethanol baths. TUNEL stainings were performed as IF experiments with the TMRRed *In situ* dell death detection kit (Roche).

### Isolation of primary cardiomyocytes and treatment with RA and BMS493

E18.5 hearts were dissected, minced and then digested in DMEM + Trypsin (100 mg/ml) for three times 15 minutes at 37°C with shaking. After each 15-minute incubation, the supernatant was removed and 5 ml FBS was added to stop the reaction. After the digestion, all solutions were pooled, run through a 70 μM filter and then spun down for 5 minutes at 1600 rpm at 4°C. The supernatant was then removed, and the pellet resuspended in DMEM + 8% FBS. The cells were then plated for 1-2 hours on uncoated plastic wells of 6 well plates for fibroblasts to adhere. The non-adherent cells containing the cardiomyocytes were then resuspended and plated on collagen coated (50 μg/ml) wells of 6 well plates. The next day the media was changed and the cardiomyocytes grown to 50-60% confluency before being treated. For RA (SIGMA) and BMS493 (TOCRIS) treatments, solutions were pre-diluted in 100% ethanol before being added directly to the media at final concentrations of 100 nM (RA for RNA-seq) or 1 μM (BMS493 and RA for qPCR analysis).

### RT-qPCR

RNA was extracted from the bottom halves of infarcted or sham adult hearts and primary cardiomyocytes using TRIzol® reagent (Invitrogen), following the manufacturer’s instructions. Reverse transcription was performed using the M-MLV reverse transcriptase in combination with oligo (dT) primers (Invitrogen). The cDNA was used as a template for quantitative PCR analysis using the SybrGREEN® Master Kit (Roche) and a Light Cycler 1.5® (Roche). Primers used for the analysis are shown in Table S3. The expression levels were normalized to *Gapdh*. For each litter or experiment ddCt values were normalized to one control dCt rather than the mean of control delta Cts. Primers (see primer table) were designed on the Universal Probe Library website (Roche).

### RNA Sequencing Analysis

TRIzol® RNA extraction was performed on four biological replicates of primary E18.5 cardiomyocytes treated with 100 nM atRA or DMSO as a control. Libraries were prepared on a Beckman Fxp Automation system, using the Illumina TruSeq stranded polyA chemistry kit. For each sample 500 ng was used as input for library preparation. Single end sequencing with an average of 20 million reads per sample was performed with the Illumina HiSeq 2000 at the EMBL sequencing center (Heidelberg, Germany). Sequences were aligned to the mouse mm10 reference genome with Burrows-Wheeler Aligner (bwa) version 0.7.12-r104 using standard parameters. Differential analysis of gene expression was calculated with the DESeq2 program from proprietary Genomatix software with a cutoff value of p<0.05 (see Table S1). The raw data files have been submitted to GEO database.

### Quantification of collagen and IF stainings on sections

For collagen staining quantification, the infarct areas from 7 sections per heart stained with Sirius Red were measured with ImageJ software and divided by the total area of the left ventricular wall. For ALDH1A2 and phospho-ERK1/2 staining levels, the positive areas for 5 sections per heart were measured with ImageJ software and divided by the area of the left ventricular wall. For TUNEL and BrdU stainings, the number of positive cells were calculated for 5 sections per heart with ImageJ software, and then divided by the total number of cells (DAPI^+^) in the left ventricular wall. For all analyses, sections were on average spaced by 40 μM, covering a total area of at least 250-400 μM.

### Statistical Analyses

Statistical analyses were performed according to the two-tailed unpaired Student t-test using Graphpad software, *p<0.05 **p<0.01, ***p<0.001. Error estimates are expressed as standard error of mean (SEM). Details of the statistical analyses and the strategies used for quantification can be found in the figure legends. The letter “n” refers to the number of individual samples/hearts (embryonic dissections and myocardial infarctions) or the number of wells (*in vitro* experiments).

## ACKNOWLEDGEMENTS

We would like to thank the staff of the animal facility and PRISM imaging platform at the iBV for their help. This work was supported by grants from the Fondation du France, ARC (SL22020605297), the ANR (ANR-11-LABX-0028-01) and La Ligue contre le Cancer (Equipe labelisée). F.D.S. and A.S. designed the project. F.D.S., A.T. and L.H.W carried out experiments. F.J.M performed all of the surgeries. The RARE transgenic construct was kindly provided by P.D. K.D.W. and A.S.R. provided critical input for experimental design and data analysis. P.D. and N.B.G. provided mice and advice. F.D.S. and A.S. wrote the manuscript, and all authors provided editorial input. The authors declare no competing interests. All data needed to evaluate the conclusions in the paper are present in the paper and/or the Supplementary Materials. Additional data are available from authors upon request.

## SUPPLEMENTARY MATERIALS

**Fig. S1:**
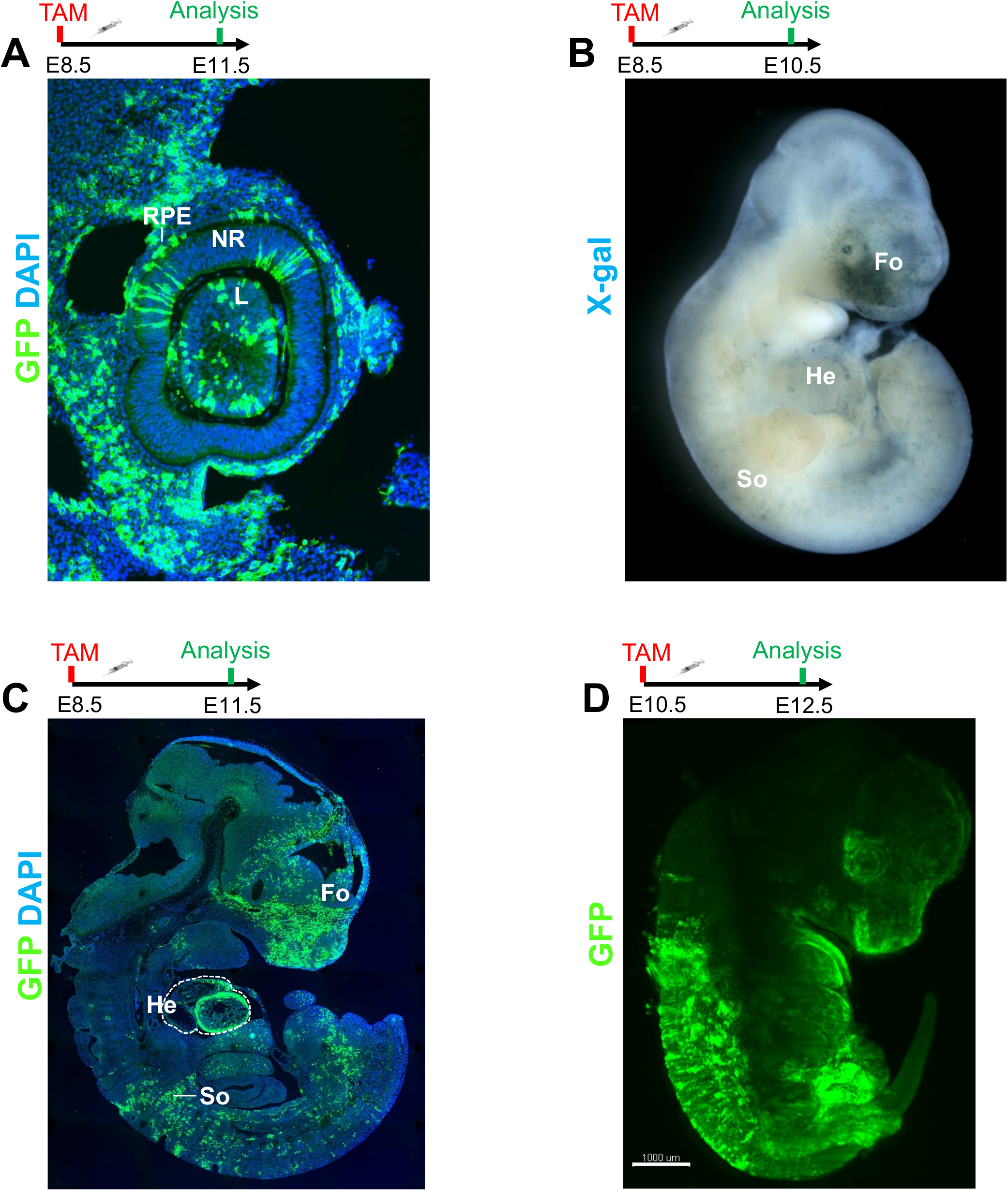
The *RARECreER*^*T2*^ line points to a reduced response at later time-points in embryonic development. **(A)** GFP IF on a *RARECreER*^*T2*^; *mTmG* embryo pulsed with tamoxifen at E8.5 and sacrificed at E11.5 reveals partial labelling of the neural retina (NR), lens (L) and retinal pigment epithelium (RPE) cells in the developing eye. **(B)** Whole-mount X-gal staining of a *RARECreER*^*T2*^; *R26L* embryo pulsed at E8.5 and sacrificed at E10.5 reveals minor labelling of segments of the forebrain (Fo), heart (He) and somites (So). **(C)** GFP IF on a sagittal section of a *RARECreER*^*T2*^; *mTmG* embryo pulsed at E8.5 and analyzed at E11.5. Recombination is detected in the forebrain, heart and a subset of somites. **(D)** Light sheet microscopy (maximal projection) of a GFP immunostained *RARECreER*^*T2*^; *mTmG* embryo pulsed at E10.5 and analyzed at E12.5 (Scale bar 1000 μM).

**Fig. S2:**
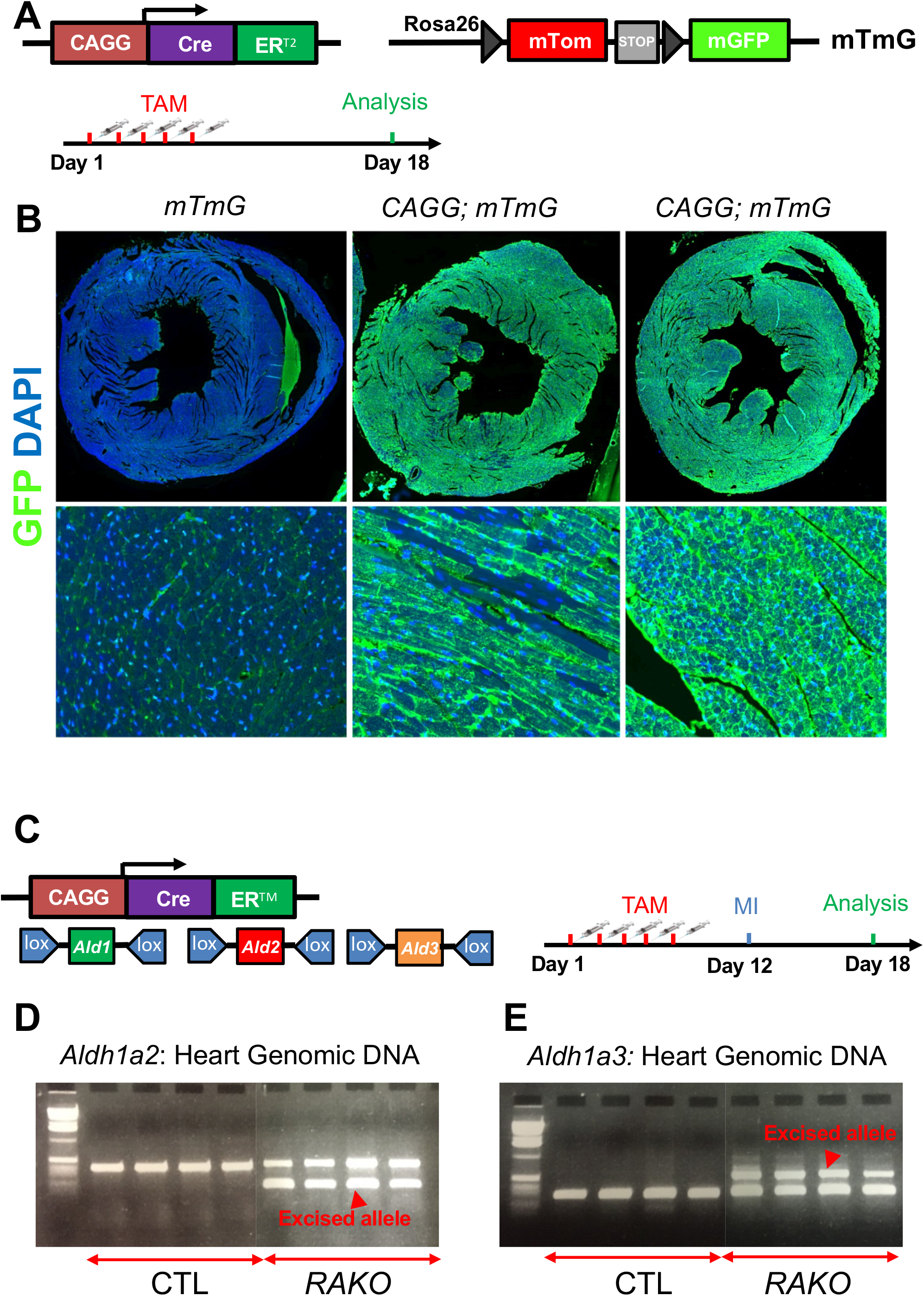
The *RARECreER*^*T2*^ line is active in the venous pole of the heart during early stages of development and in the myocardium during later stages, and myocardial labeling during mid-gestation is detected with the *RARECreER*^*T2*^ line B. **(A)** X-gal and Eosin staining on sections of a *RARECreER*^*T2*^ *R26L* heart pulsed at E6.5 and sacrificed at E9.5 (from wholemount stainings in Fig. 1). AVC = atrioventricular canal, OFT = outflow tract. **(B)** X-gal and Eosin staining on sections of a *RARECreER*^*T2*^ *R26L* heart pulsed at E10.5 and sacrificed at E13.5 (from wholemount stainings in Fig. 1). My = myocardium, Ep = epicardium. **(C)** Administration of tamoxifen at E10.5 to embryos from a second *RARECreER*^*T2*^ line (line B) crossed with the mTmG line demonstrates a strong GFP response in the compact myocardium when analyzed at E14.5 (n=3 embryos analyzed). TRO = Troponin T. Scale bars mosaics: 100 μM, close ups 40 μM.

**Fig. S3:**
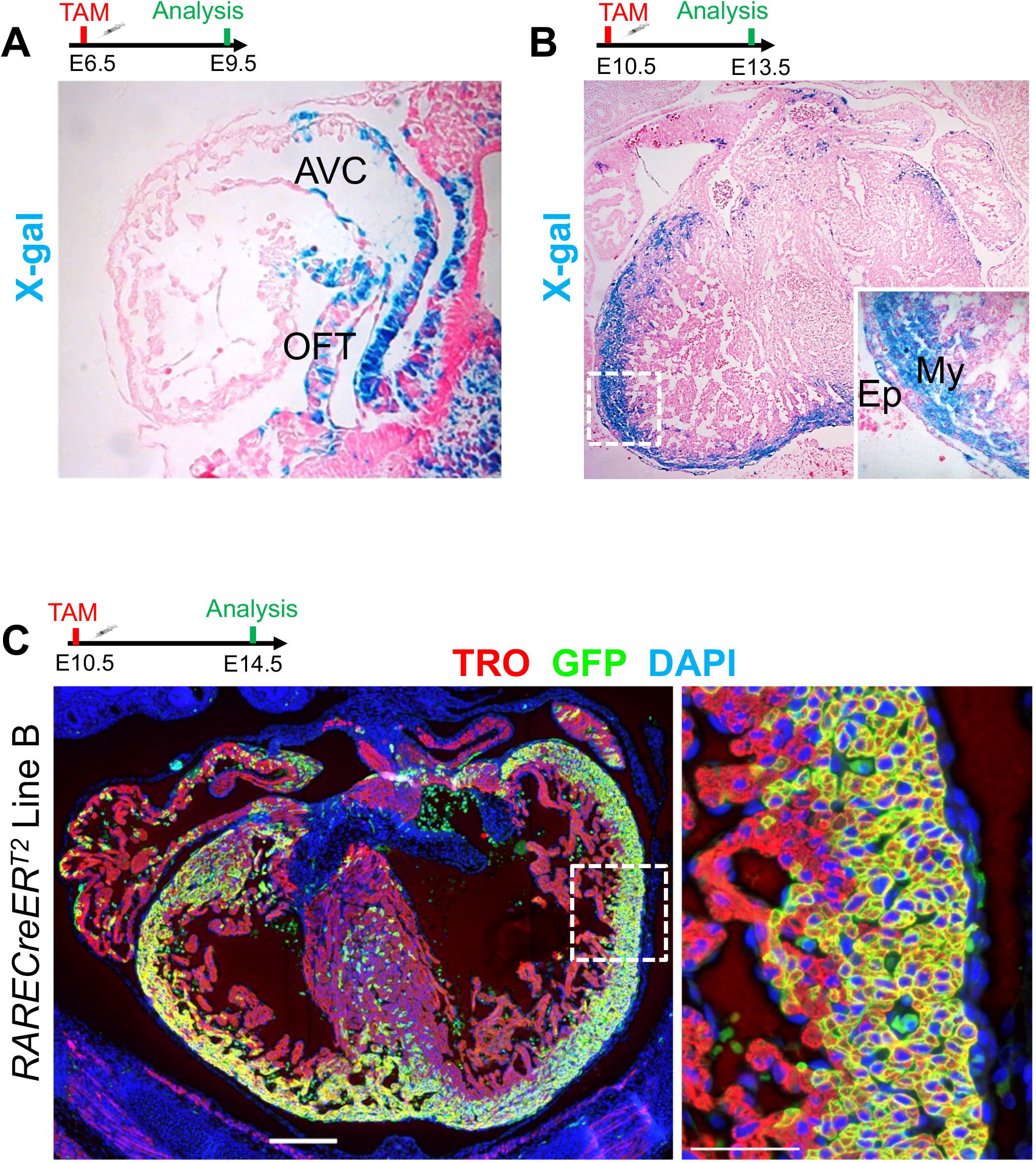
The *CAGGCreER*^*TM*^ line is highly active in adult hearts but leads to incomplete excision of *Raldh* floxed alleles. **(A)** Schematic illustrating strategy used to test the recombination efficiency of the *CAGGCreER*^*TM*^ line with the *mTmG* reporter allele. Tamoxifen was administered 5 times to *CAGGCreER*^*TM*^:*mTmG* adult males and hearts were analyzed 13 days after the final injection. Grey arrowheads represent *loxP* sites. **(B)** GFP IF on *CAGGCreER*^*TM*^:*mTmG* hearts reveals very efficient recombination in nearly all cell types. Two representative *CAGGCreER*^*TM*^:*mTmG* hearts shown. **(C)** Schematic of strategy to generate *Aldh1a1,2,3 (Ald1,2,3)* KO adult mice (referred to as *RAKO*s) followed by MI surgery. **(D-E)** PCR analysis of genomic heart DNA from *RAKO* adults subjected to MI reveals excision of floxed *Aldh1a2* **(D)** and *Aldh1a3* **(E)** alleles (red arrowheads). Amplification of the non-excised alleles still occurs suggesting incomplete recombination in *RAKO* hearts.

**Fig. S4:**
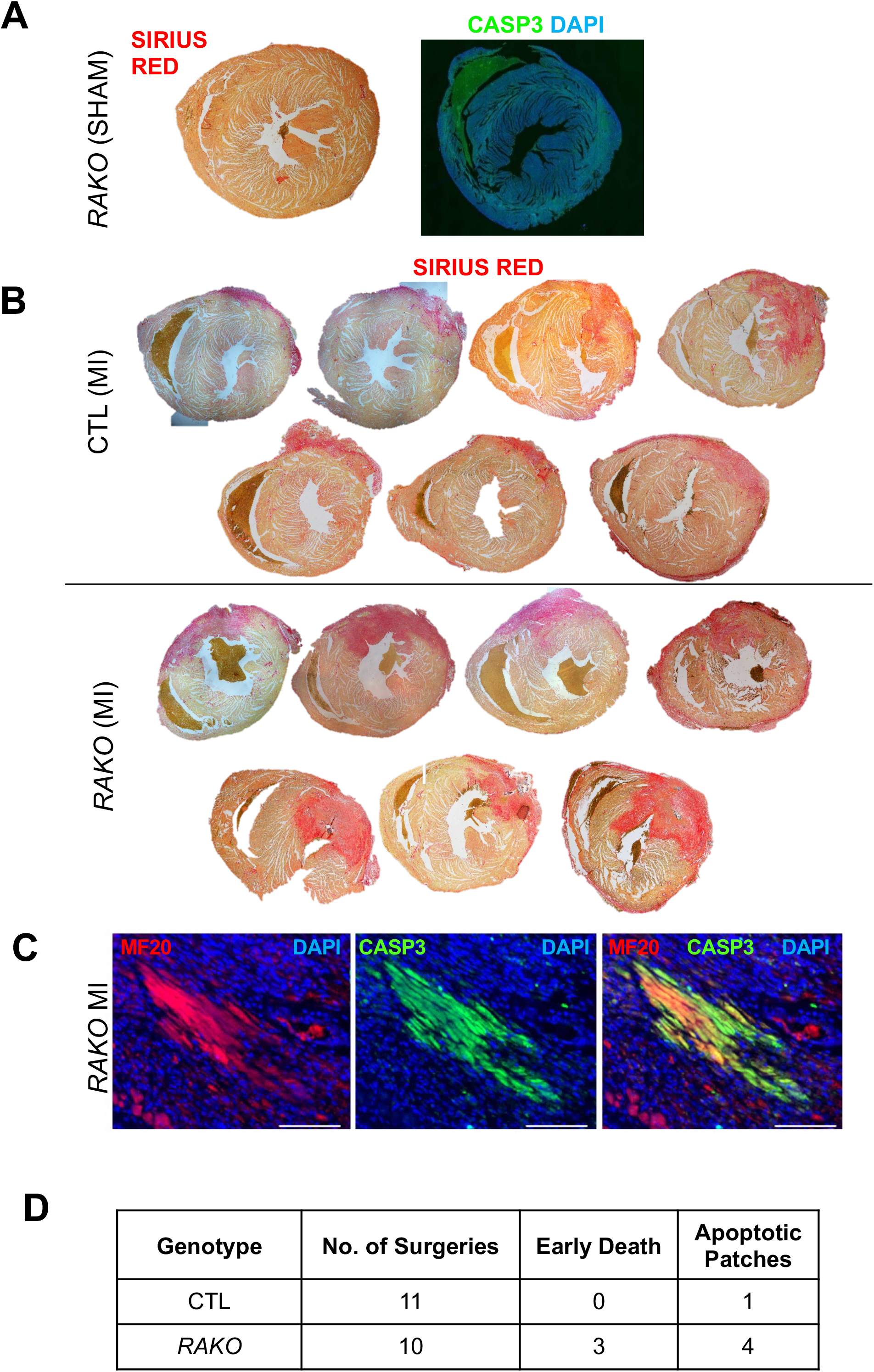
Increased infarct zones in *RAKO* hearts subjected to myocardial infarction. **(A)** *RAKO* sham hearts do not display any adverse remodeling defects or apoptosis as shown by Sirius red staining and active caspase 3 (CASP3) IF. **(B)** Sirius red staining of various MI hearts reveals consistently larger infarct sizes in *RAKO* hearts when compared to control (CTL) hearts. For each heart representative image from largest portion of infarct shown. **(C)** Apoptotic patches observed in *RAKO* mice after MI are cardiomyocytes as revealed by co-IF for active caspase 3 and MF20. **(D)** Table showing *RAKO* MI hearts exhibit a higher incidence of early death (prior to analysis at 6 days post MI) and more “apoptotic patches”.

**Fig. S5:**
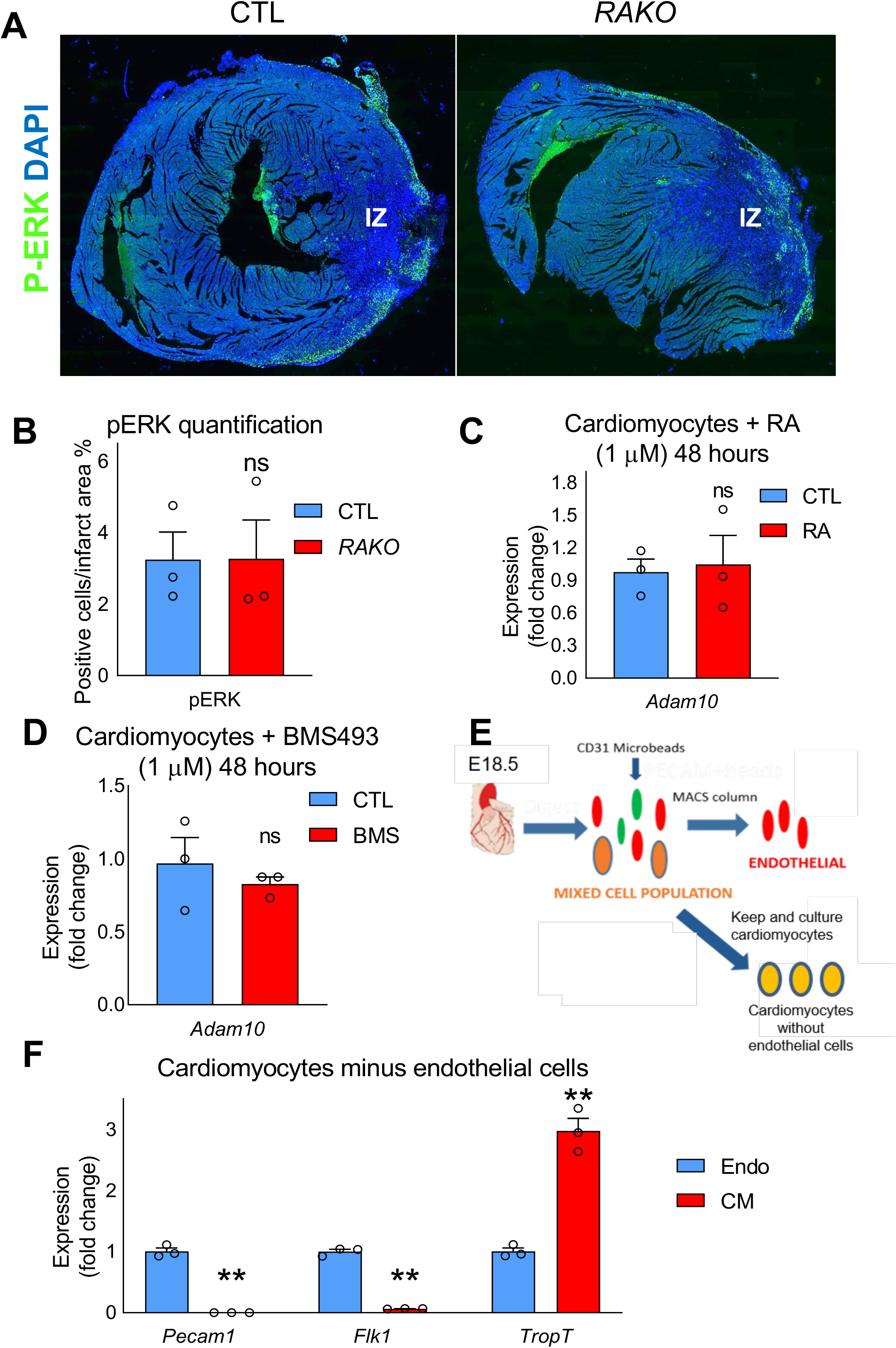
MAP kinase signaling is not significantly altered in RAKO infarct hearts. **(A)** IF against phospho-ERK1/2 reveals no major differences in *RAKO* MI hearts when compared to controls. **(B)** Quantification of phospho-ERK1/2 levels reveals no significant differences between *RAKO* and control hearts. Phospho-ERK1/2 pixel area was measured using ImageJ software and was normalized to total infarct area. Columns are means ± SEM (n=3 hearts). **(C)** qPCR analysis of primary cardiomyocytes treated with RA (1 μM) for 48 hours demonstrates no significant difference in *Adam10* expression. Data are expressed as fold change vs controls and columns are means ± SEM (n=3). **(D)** qPCR analysis of primary cardiomyocytes treated with the RAR reverse agonist BMS493 (1 μM) for 48 hours demonstrates no significant difference in *Adam10* expression. Data are expressed as fold change vs controls and columns are means ± SEM (n=3). **(E)** Schematic showing strategy to remove endothelial cells from primary cardiomyocyte cultures using CD31-magnetic beads. MACS = magnetic activated cell sorting. **(F)** qPCR analysis of cell isolation from **(E)** reveals purified cardiomyocytes (CM) express very low levels of endothelial (Endo) markers (*Pecam, Flk1*) and high levels of Troponin T (n=3). Data are expressed as fold change vs controls and columns are means ± SEM. PERK-phospho-ERK, TropT = Troponin T. All statistics two tailed t-test assuming unequal variance, **p<0.01, ns = not significant.

**Fig. S6:**
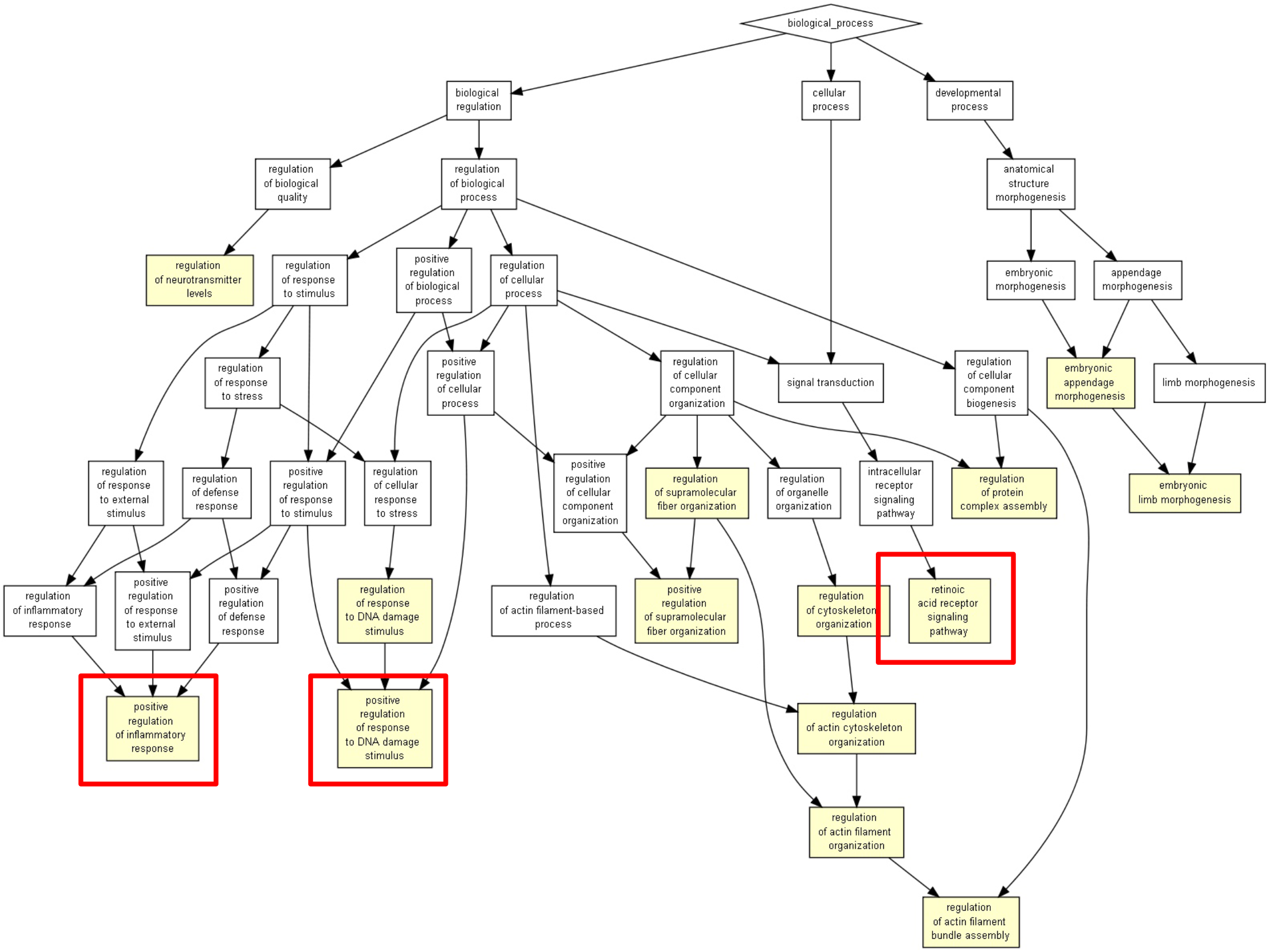
Gene ontology analysis of RNA sequencing data. **(A)** Gene ontology analysis of biological processes altered in RA-treated primary cardiomyocytes using RNA sequencing data from top 500 deregulated genes. Analysis performed with Gorilla software.

**Supplementary Table 1:** Differential expression analysis (RNA Seq) on E18.5 primary cardiomyocyte cultures treated with all-trans retinoic acid. (see Excel file)

**Supplementary movie 1:** 3D analysis of *RARECreER*^*T2*^ embryos crossed with the *mTmG* reporter, pulsed with tamoxifen at E7.5 and sacrificed at E10.5. Staining was performed with antibodies against GFP (green). Heart and mesonephros were highlighted using antibodies against GATA4 (magenta) and PAX2 (white), respectively.

**Table S2:** List of Antibodies used in this study

| **Protein** | **Host** | **Type** | **Dilution** | **Manufacturer** | **Catalogue No.** |
| --- | --- | --- | --- | --- | --- |
| **GFP** | Chicken | polyclonal | 1:400 | Abcam | ab13970 |
| **ALDH1A2** | Rabbit | polyclonal | 1:300 | Sigma | ABN420 |
| **Myosin D** | mouse | monoclonal | 1:200 | Santa Cruz | sc32758 |
| **MF20** | mouse | monoclonal | 1:20 | DSHB | AB 2147781 |
| **αSMA** | mouse | monoclonal | 1:1000 | Santa Cruz | sc53015 |
| **WT1** | mouse | monoclonal | 1:100 | Dako | M3561 |
| **Vimentin** | Chicken | Abcam | 1:1000 | Abcam | ab24525 |
| **Active caspase 3** | Rabbit | polyclonal | 1:200 | R & D | AF835 |
| **Phospho-ERK1/2** | Rabbit | polyclonal | 1:400 | Cell Signaling | 4370 |
| **Troponin T** | Mouse | monoclonal | 1:300 | Invitrogen | MA5-12960 |
| **GATA4** | Goat | polyclonal | 1:500 | Santa Cruz | sc1237 |
| **PECAM** | Goat | polyclonal | 1:200 | Santa Cruz | sc1506 |
| **Transgelin (SM22α)** | Rabbit | polyclonal | 1:400 | Abcam | ab14106 |

**Table S3:**
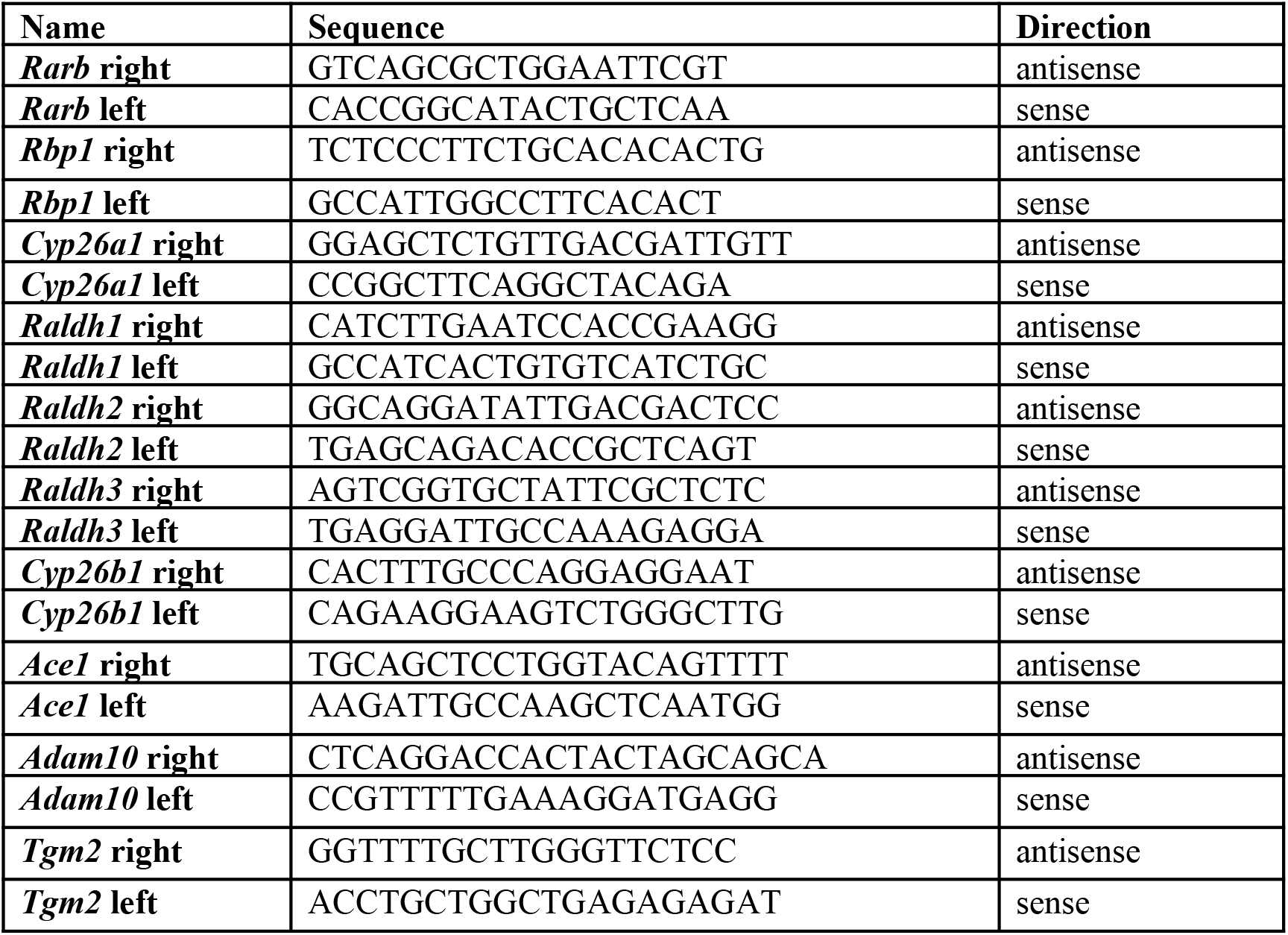
List of primers used for qPCR analysis in this study

